# Of Shared Homes and Pathways: Free-Ranging Dog Movement and Habitat Use in a Human-Wildlife Landscape in India

**DOI:** 10.1101/2025.06.30.662475

**Authors:** Sanjana Vadakke Kuruppath, Nilanjan Chatterjee, Ramesh Krishnamurthy

**Affiliations:** #12, Valmiki Nagar, Thiruvanmiyur, Chennai 600041, India; Senckenberg Biodiversity and Climate Research Institute, Frankfurt am Main, 60325, Germany; Wildlife Institute of India, Dehradun, Uttarakhand 248001, India

**Keywords:** Human-wildlife conflict, movement ecology, human shield hypothesis, step- selection analysis

## Abstract

**Background:** Free-ranging dogs are widely considered to have negative impacts on wildlife in shared landscapes. Understanding their use of these landscapes is therefore essential to develop effective management strategies in wildlife-adjacent regions. In this study, we investigated the movement patterns of free-ranging dogs in the protected area matrix of Mudumalai Tiger Reserve (MTR) in Southern India.

**Methods:** Fifteen owned free-ranging dogs (FRD) were collared from two adjacent villages with different levels of human habitation. We hypothesized that increased settlement cover would provide dogs with a greater human shield effect, allowing them more freedom of movement. We estimated dog activity ranges and movement patterns using autocorrelated kernel density estimates (AKDE). Large-scale movement was characterized through multiple metrics (intensity of space use and mean distance from home) and compared across villages to understand underlying drivers. Lastly, we quantified fine-scale movement and habitat selection using integrated step selection analysis (iSSA).

**Results:** We found that FRD in MTR had directionally dependent home-ranging behavior and small activity ranges, with the mean AKDE activity range being 9.88 ± 7.69 ha (median = 6.09 ha, range 3.28-26.16 ha), and primarily utilized the area within 500 m of their homes. None of the movement metrics varied significantly between villages, except for intensity of use, which was higher in one village than the other, suggesting that dogs in the less populated village perceived a greater threat from surrounding wildlife and were more driven to seek refuge inside their activity ranges. iSSA revealed that dogs selected strongly for human settled or human-modified habitats, and moved significantly faster in forest land than any other habitat.

**Conclusions:** Our findings indicate that owned FRD pose a relatively lower threat to wildlife in MTR than feral dogs may. They also support the human shield hypothesis, show that FRD display behavioral plasticity at fine scales of <5km, and highlight the role of human land use intensity in shaping the movement of domestic dogs. As humans can thus mediate potential effects of dogs on wildlife, anthropogenic factors should be taken in consideration when designing management strategies that aim to curb dog movement.

## Introduction

As the most abundant carnivore on earth with a long history of domestication [1], domestic dogs (*Canis familiaris*) are almost as widely distributed as humans across the world [1], with rural dogs forming a sizable category of the population in underdeveloped areas [2]. These dogs are highly likely to be free-ranging, with previous studies on rural dog demography across the world indicating that a major proportion of rural dogs are free to roam without any restraint, whether alone or in the company of humans [3–6]. In areas close to natural ecosystems, free-ranging dogs (FRD) can have a range of negative impacts on wildlife [7,8], ranging from mild to severe depending on the local socio-ecological context [9]. Broadly, the most common negative impact of domestic dogs on threatened wildlife is predation, estimated at 78.9%, followed by disturbance (such as barking, chasing or harassment) and disease transmission at 7.0% and 4.5% respectively, and lastly competition with native carnivores (1.5%) and hybridization with wild canids (1.0%) [10]. While the intensity of these impacts is unclear, there is no doubt that they are globally pervasive [10].

In order to clarify the true extent of dog-related interference with wildlife, understanding how FRD utilize wildlife-adjacent landscapes is a necessary avenue of research, as patterns of movement into wildlife habitat underpin all subsequent impacts on wild species. These patterns can vary widely across ecosystems, with a metric as fundamental as the home range (or the activity range, for short-term studies) varying regionally, seasonally, or individually, as previous studies from Australia [11–16], Chile [17], Kenya [18], Cambodia [19], and Mexico [20,21] have shown. Therefore, aspects of dog movement ecology that are of particular interest from a wildlife conservation perspective, such as ranging behavior and habitat selection, may be equally varied and may be influenced by a range of natural as well as anthropogenic factors. Indeed, given the highly social and commensal relationship between dogs and humans [1], even subtle anthropogenic factors may significantly influence dog movement and behavior [10]. For example, dogs that disrupted sea turtle nests to scavenge for eggs in Mexico were fed significantly less food by their owners than to dogs that did not scavenge [21].

While anthropogenic factors are usually considered to be resources that humans actively provide dogs with, such as food, there may also be passive benefits to existing in close proximity to humans. The human shield effect, where prey species are hypothesized to seek refuge from predators that fear humans by increasing their use of human-modified areas [22], is an example of one such potential benefit. Although this hypothesis was developed with reference to the behavior of wild prey species [22], a diverse range of apex carnivores have also been documented to prey on dogs across the world [23]. There is, therefore, scope for FRD to also benefit from the protection offered by human presence, by using settlement areas to avoid encountering predators. As a result, different levels of habitation could affect how freely dogs utilize landscapes shared with wildlife, providing new insight into how FRD patterns of movement differ across anthropogenic contexts.

The need to elucidate such potential patterns underlying dog movement ecology is particularly pressing in India, where the majority of the dog population is free ranging [24] and may often come into contact with increasingly fragmented natural ecosystems [25] that harbor a rich diversity of wildlife [26]. Previous studies have investigated other impacts of dogs on wildlife in India [27–29], with one study recording reports of negative interactions between dogs and 80 wild species [30]; however, to our knowledge, only two studies have estimated FRD movement ecology in rural or wildlife adjacent areas, both in the state of Maharashtra [31,32], of which only Vanak and Gompper (2010), utilized VHF telemetry to estimate home range and habitat use. While non-invasive methods such as camera trapping can be used to estimate movement parameters, they are relatively time consuming, effort-intensive [33], and can be spatially constrained compared to telemetry (specifically GPS telemetry), which allows for the collection of individual-level, precise, fine-scale data around the clock across large areas [34]. When studying FRD that may exhibit highly context-dependent spatial behavior, such data become particularly valuable.

We address this research gap with a fine-scale GPS telemetry study on rural FRD from the multiple-use landscape of a Tiger Reserve in Southern India, where we investigate how FRD movement and utilization of multiple habitat types are affected by different levels of anthropogenic land use intensity. We collared FRD in two different villages, both situated inside the Reserve boundary and zone of influence, with varying levels of human population, built-up area, and village-forest boundary length. We hypothesized that while FRD would primarily use settlement areas, a higher proportion of built-up land, in conjunction with a smaller village-forest boundary, would produce a human shield effect to allow dogs more freedom of movement in a forested landscape harboring large carnivore species. To test this hypothesis, we designed a study with two objectives: 1. Estimate and compare FRD activity ranges and movement metrics between villages and 2. Evaluate how FRD use of habitat types differs between villages.

## Materials and Methods

### Study area

The study was conducted in the villages of Masinagudi and Moyar, in Mudumalai Tiger Reserve (MTR), Tamil Nadu, India (Figure 1). MTR is a protected area contiguous with multiple other wildlife sanctuaries, forming an integral part of the Nilgiri Biosphere Reserve in the highly biodiverse Western Ghats. Several villages are embedded within MTR, forming a matrix landscape where human settlements and major roads are interspersed within the forest. The study sites, Masinagudi and Moyar, are situated towards the center of the Reserve on the Sigur Plateau, where the primary vegetation types are dry deciduous and dry thorn forest. MTR supports a variety of mammalian wildlife, including tiger (*Panthera tigris*), leopard (*Panthera pardus*), elephant (*Elephas maximus*), dhole (*Cuon alpinus*), chital (*Axis axis*), sambar (*Rusa unicolor*), blackbuck (*Antilope cervicapra*), gaur (*Bos gaurus*), and sloth bear (*Melursus ursinus*) [35]. Leopards in particular have been documented to prey on dogs in India since colonial times [23] and commonly do so in MTR at present (SVK, unpub. data).

**Figure 1:**
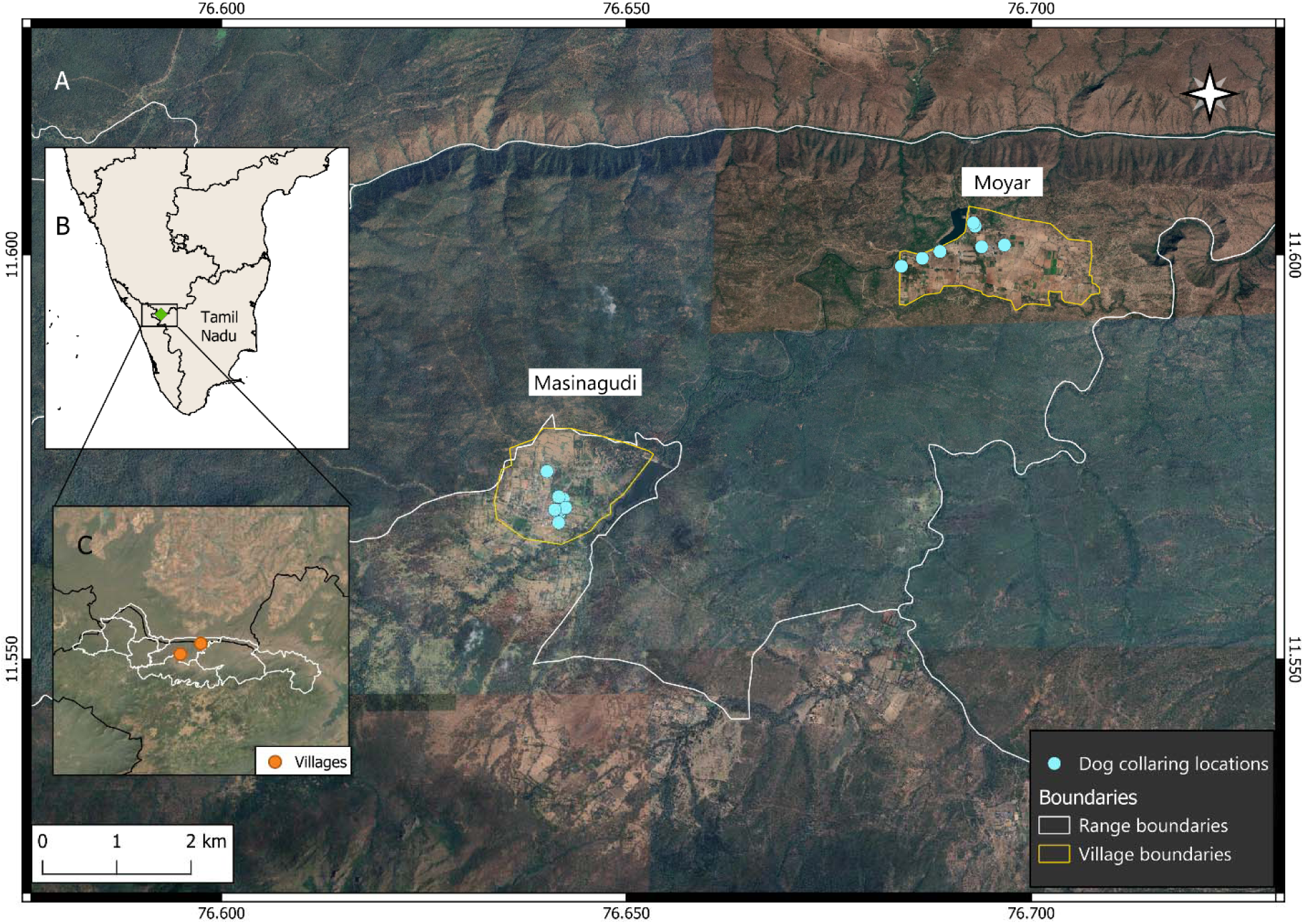
Map of the study area, showing dog collaring locations inside the villages of Masinagudi and Moyar (A) inside MTR (B) in peninsular India (C). Collaring locations are shown for the dogs whose data were included in this study.

Masinagudi is smaller in area (2.29 km^2^) with a larger population of 1641 households (as of the 2011 Government of India census; the current number is likely higher), has more dense and contiguous built-up area, and has the most developed infrastructure of all villages on the Sigur Plateau. A major source of income for most Masinagudi residents is from tourism, by running homestays or driving safari jeeps. In contrast, Moyar is slightly larger (2.37 km^2^) but hosts only 486 households, with houses clustered in a few small settlements separated by stretches of fallow or agricultural land, which occupy the majority of the village area. Here, income is primarily derived either from agriculture or from working at a hydroelectric project at the Moyar dam. As the villages are situated within the forest, village-forest boundaries are routinely utilized by humans and domestic animals as well as wildlife in both places, although several wild species (including spotted deer (chital), sambar, wild pig, elephant, and leopard) enter Moyar more frequently than Masinagudi (SVK, unpub. data). Masinagudi and Moyar are situated 4.7 km apart on the Sigur Plateau, connected by a 6.8 km long road.

Most people commonly keep dogs in this landscape, primarily as security against wildlife but also as pets (SVK, unpub. data). Dog reproductive management has been ongoing since 1997 in the form of vaccination and sterilization programs, carried out by local animal welfare organizations (India Project for Animals and Nature, IPAN, and Worldwide Veterinary Service- India, WVS-I). At present, almost 70% of dogs are vaccinated and sterilized, and the male:female ratio is almost 1:1 (SVK, unpub. data).

### Collar deployment

GSM GPS collars (by Arcturus Inc., Bengaluru, India) were deployed on owned dogs with the permission and assistance of the owners. These collars relied on mobile 2G network to transmit collected data to a data server. A total of 26 dogs were collared between September-November, 2024. The collars were set to record a location every 5 minutes, with an accuracy (measured by horizontal dilution of precision, or HDOP) of ten meters or less, and we aimed to collect data for maximum of ten days for each dog. Ten days was thought to be a reasonable duration, as other studies have tracked dogs for shorter periods [11,18,36]. Additionally, residents were aware that the Forest Department prohibits the presence of dogs in the forest and were afraid that allowing their dogs to be collared would result in punitive action, making it difficult to request that collars be kept on their dogs for longer. Indeed, in practice, the number of deployment days varied as owners sometimes requested that the collar be removed early, the collar was removed without our knowledge, or the collar was reported to have fallen off on its own, resulting in a range of deployment durations of 3-10 days.

We selected adult FRD at random from households in several different localities within each village in order to obtain a diverse sample pool, although we attempted to maintain an even sex ratio. Collaring was accomplished after the owner gave permission and in many cases the collar was placed by the owner under supervision of the collaring team. Most collared FRD belonged to people living and working in the interior of the village, as dog owners living near the village-forest boundary were especially reluctant to allow their dogs to be collared. Owners also reported that their dogs sometimes chose to stay near the house or roam on their own, rather than accompany them during the day. Therefore, while dogs in MTR generally go with their owners to agricultural fields or during livestock grazing, we did not expect collared FRD to closely mirror human movement patterns, and no collared dog was expected to routinely visit the forest in the company of their owner. In particular, it should be noted that collared FRD presence in the forest would not reflect associated human movement for hunting; apart from the danger posed to humans by elephants, sloth bears and large carnivores, hunting is strictly illegal in protected areas in India. Therefore, even if people did engage in any kind of poaching, they would be highly unlikely to do so regularly, and would not take a collared FRD with them.

After collaring, the dogs were photographed and several demographic variables were recorded using the mobile phone application Epicollect [37], including sex, sterilization status, approximate age according to the owner, and the GPS location of the owner’s home. All dogs were adults and were fed a minimum of once and a maximum of three times per day by the owner.

## Analysis

We collated the GPS data that was successfully collected during deployment and uploaded them to Movebank, a free online repository for animal movement data from across the world [38].

Outliers were removed through the online Movebank interface in two steps, first using a speed filter (excluding points that would require the dog to move at a speed > 5 m/s) and second manually. 5 m/s was considered a lenient upper limit, since while dogs are capable of reaching this speed, they would be unlikely to maintain it consistently for the five-minute interval between fixes to run for a stretch of 1.5 km. We considered that speeds below this limit would account for any potential encounters with wildlife, whether chasing prey species or escaping from predator species.

Points removed manually were implausible for accessibility reasons (e.g., the middle of a fenced waterbody) or distance reasons (e.g., random single points > 500 m away from clustered preceding and subsequent points), and were presumably due to poor satellite reception.

Subsequent data analysis was conducted with the data downloaded from Movebank (see Data Availability) in order to ensure conformity to a standard format. Dogs with excessively patchy data (fewer than 300 points collected) were excluded, since these constituted approximately only one day of data and they often had gaps of multiple days in between fixes, making it unfeasible to use them for a robust analysis. Lastly, points with an accuracy (HDOP) of less than 10 m were removed from the dataset.

### Movement Metrics

To examine the difference in dog movement patterns between villages, we utilized multiple different movement metrics, namely the activity range, the intensity of use, and the mean distance between the owner’s home and each GPS fix for each dog. We estimated the activity range for each dog, or the area traversed by the dog during the short-term period of deployment [19] with the package *ctmm* (v1.2.0) [39] in R version 4.4.1[40], using autocorrelated kernel density estimation (AKDE), similar to Wilson-Aggarwal et. al, 2021. Six models were fitted to each dog’s spatial data, and the best fit model was selected using the *ctmm.select* function, which ranks models according to the Aikake Information Criterion [39]. The models included Ornstein-Uhlenbeck (OU), OUF (Ornstein-Uhlenbeck with foraging), and IID (Independent and Identically Distributed), of which IID alone does not consider autocorrelation. After model selection, we estimated the 95% AKDE activity range for the dog based on the best fit model using the *akde* function [39].

The second movement metric, the intensity of use, is a measure of the active time spent by the animal per unit area, defined as the ratio of the total distance moved during the sampling period to the square root of the 100% minimum convex polygon (the maximum area used by the animal) [41,42]. We estimated it with the package *amt* [43], using the *intensity_use* function. The third movement metric was the mean distance between the owner’s home and each GPS fix for each dog. We calculated the distance between home and the dog’s location using the package *sf* [44] for every set of GPS coordinates recorded, and subsequently averaged the distances to provide the mean as a measure of how far away from home each dog tended to travel. We repeated this process with the distance from the centroid of the activity range to assess whether there was any significant difference between the two.

We further categorized the movement data with respect to day and night for the mean distance from home, and with respect to sex of the dog and the village of origin for all three movement metrics. We hypothesized that FRD would move more during the day to avoid wild animals at night, that a higher amount of built-up area and a smaller village-forest boundary in Masinagudi would allow FRD more freedom of movement than in Moyar, and that males would cover larger areas that than females despite both being sterilized, as has been recorded elsewhere [e.g. 45].

Overall, since these FRD were owned and provisioned, we expected that they would tend to remain in the vicinity of their owners’ homes for most of the time they were active. The level of human subsidy was not included as a variable in this study since feeding was regular. As the collaring period was relatively short, the difference between sunrise and sunset times during the study was negligible; therefore, we used 0600-1800 hours as day time and 1800-0600 hours as night. We tested the difference between movement metrics with respect to these variables using the non-parametric Wilcoxon rank sum test.

We also estimated the dog daily activity pattern using the nonparametric kernel density estimation method, expecting FRD to occupy a strongly diurnal diel niche. GPS data from individual dogs were used to calculate movement rates as the distance between successive locations divided by the time between them. We then converted the rates to randomly distributed pseudo-detections within the time interval between GPS fixes, so that a higher speed translates to more pseudo-detections, to obtain a general, continuous activity pattern [46].

### Habitat Selection

To evaluate the selection and movement of the dogs in different habitats, we first classified the areas of each village into settlement, agriculture, fallow or scrubland (uncultivated land beyond settlements but within village boundaries), and forest land (Fig. 2). Forest land included both deciduous forest and thorn forest vegetation types. We used land-use/cover Sentinel-2 imagery at a resolution of 10 meters, classifying specific habitat types (such as fallow/scrub land) manually, given the fine scale of the study area. As an initial coarse assay of habitat selection, we used landscape imagery of approximately 70 sq. km, enough to maintain a buffer of 1 km in all directions from both villages, and calculated the proportion of dog presence with respect to habitat availability for each dog.

**Figure 2.**
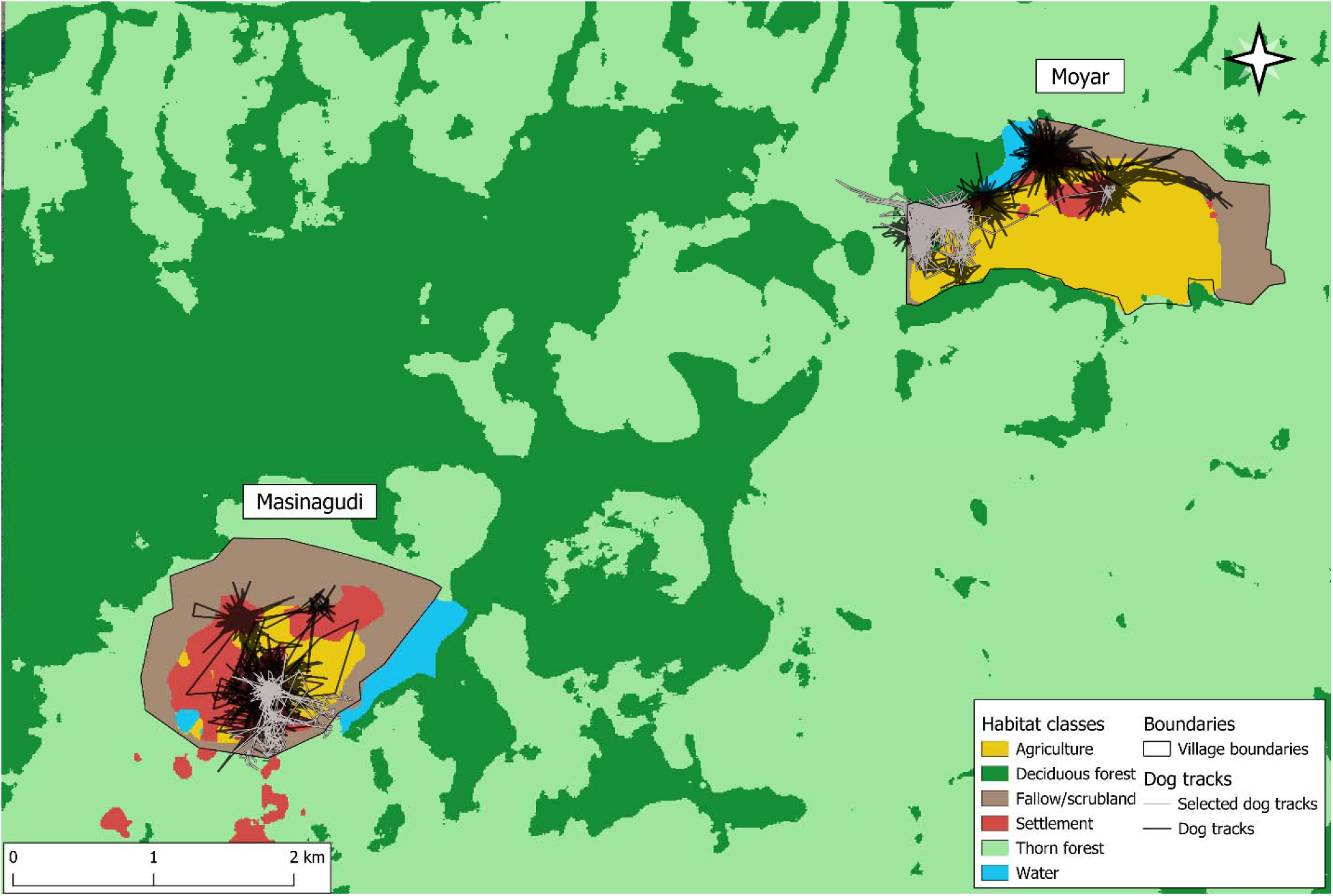
Land use/cover classes of the study area. Black lines indicate dog movement tracks in the vicinity of each village. One track is highlighted in grey in each village to exemplify the dogs with the largest activity ranges from each village.

We then utilized integrated step-selection analysis, or iSSA [47] to study the fine-scale influence of habitat covariates on the habitat selection and movement rate of the dogs. iSSAs contrast the environmental attributes of each animal location with a set of random locations generated from the distribution of movement parameters (step lengths and turn angles) of the steps the dogs took [47]. As with other habitat selection analyses, iSSAs compare covariate values at used (animal GPS locations) and random points (locations generated using the distribution of the movement parameter) for each animal step. iSSAs are considered to be an improved approach to habitat select analysis as the framework can simultaneously estimate coefficients of habitat selection and movement rate, potentially increasing the accuracy of both [47,48]. We generated 20 available steps per used step from the observed movement parameter distribution of the dogs. As the data was collected at a very high resolution, we resampled the movement data to 15 minutes and 30 minutes to control and test iSSA models for spatial autocorrelation. The village of origin of the dogs was considered as a random effect for habitat and movement to elucidate differences in movement between the two villages. We utilized the Poisson trick [48] and the variability in movement across villages using the random effect approach [49]. Models were fitted using the package “glmmTMB” [50] and visualized using package ggplot2 [51]. All codes to reproduce the analysis are publicly available (see Data Availability).

## Results

Of the initial 26 collared dogs, the data from four dogs were completely lost due to collar error or premature collar removal by unknown persons, leaving us with data from 22 dogs totaling 27,850 GPS points. Of these, 4144 points were removed as they were null points (i.e. points recorded as 0,0 when the collar failed to acquire a GPS fix), 49 points were removed with the Movebank speed filter, 186 were removed manually, and three points were removed due to low accuracy, resulting in a final count of 23,468 points. Of the 22 dogs included in the Movebank dataset, seven were excluded during analysis as their data were few in number and excessively patchy, presumably due to satellite connection or mobile network coverage issues. The final dataset therefore comprised GPS location data from 15 FRD with robust datasets, with the number of points totaling 22277.

The set of 15 FRD were almost equally divided both in terms of sex as well as the village of origin, with 8 females and 7 males, and 8 FRD from Masinagudi and 7 FRD from Moyar. Of these, all were sterilized apart from one male dog (Table 1). The mean age of the all dogs tracked was 5 (range = 3 - 8). The mean number of tracking days was 8.07 ± 2.40 (range = 3 - 10). The mean number of GPS points per dog was 1485 ± 738 (range 619 - 2783) and post outlier filtering, the precision of the data was high (mean HDOP = 1.66 m ± 0.70 m, range = 0 - 9.13 m).

**Table 1:** Individual dog demographic parameters and tracking data. Values rounded to two decimal points.

| Dog ID number | Village of origin | Sex | Age | Sterilization status | No. of fixes | No. of days tracked |
| --- | --- | --- | --- | --- | --- | --- |
| D1 | Masinagudi | Male | 7 | Yes | 1169 | 8 |
| D6 | Masinagudi | Female | 3 | Yes | 906 | 10 |
| D8 | Masinagudi | Male | 5 | Yes | 2174 | 10 |
| D9 | Masinagudi | Female | 4 | Yes | 647 | 7 |
| D11 | Masinagudi | Female | 5 | Yes | 1140 | 8 |
| D12 | Masinagudi | Male | 3 | Yes | 1501 | 8 |
| D13 | Masinagudi | Female | 5 | Yes | 619 | 3 |
| D14 | Masinagudi | Female | 3 | Yes | 635 | 9 |
| D2 | Moyar | Female | Unknown | Yes | 2446 | 9 |
| D3 | Moyar | Male | Unknown | Yes | 2783 | 10 |
| D4 | Moyar | Male | 5 | No | 2172 | 10 |
| D5 | Moyar | Female | 7 | Yes | 1874 | 10 |
| D7 | Moyar | Female | 8 | Yes | 757 | 3 |
| D10 | Moyar | Male | 8 | Yes | 1207 | 6 |
| D15 | Moyar | Male | 5 | Yes | 2247 | 10 |

The activity range best fit model for individuals from both villages was OU anisotropic, often with a ΔAIC of approximately 2 with the next best model, which was OUF anisotropic (see Supplementary Table S3 for all ΔAIC values), while IID was ranked last for all individuals, indicating that collared FRD displayed directionally dependent home-ranging behavior. The mean AKDE activity range estimate across all dogs was 9.88 ± 7.69 ha (median = 6.09 ha, range = 3.28 ha – 26.26 ha) (Table 2). We found no significant difference between mean distance from home and mean distance from the centroid, with the distance between home and centroid sometimes varying by as little as 10 meters; therefore, all further analysis was conducted with respect to distance from home only.

**Table 2:** Individual dog movement metrics. Values rounded to two decimal points.

| Dog ID number | Village of origin | Activity range (ha) | Activity range, 95% confidence interval | Distance from home, mean (m) | Distance from home, range (m) | Intensity of use |
| --- | --- | --- | --- | --- | --- | --- |
| D1 | Masinagudi | 11.46 | 10.32 -12.66 | 76.84 | 0.24 – 667.73 | 87.84 |
| D6 | Masinagudi | 26.16 | 22.97 - 30.67 | 127.27 | 0 – 606.64 | 72.46 |
| D8 | Masinagudi | 4.56 | 4.37 - 4.83 | 45.34 | 0 – 859.40 | 137.94 |
| D9 | Masinagudi | 9.88 | 7.84 - 12.19 | 52.69 | 1.56 – 900.99 | 54.61 |
| D11 | Masinagudi | 3.81 | 3.58 - 4.05 | 39.94 | 1.09 – 341.86 | 131.48 |
| D12 | Masinagudi | 7.19 | 6.46 - 7.96 | 54.12 | 0 – 923.06 | 87.59 |
| D13 | Masinagudi | 14.40 | 10.32 - 19.12 | 79.28 | 1.56 – 690.90 | 62.86 |
| D14 | Masinagudi | 5.80 | 4.81 - 6.88 | 49.05 | 0 – 777.84 | 48.89 |
| D2 | Moyar | 4.49 | 4.24 - 4.74 | 58.95 | 1.09 – 428.27 | 213.08 |
| D3 | Moyar | 4.19 | 3.87 - 4.52 | 50.86 | 0 – 913.85 | 166.23 |
| D4 | Moyar | 23.86 | 21.32 - 26.54 | 124.93 | 0 – 1197.06 | 179.03 |
| D5 | Moyar | 3.39 | 3.22 - 3.57 | 41.51 | 1.09 – 576.37 | 146.01 |
| D7 | Moyar | 3.28 | 3.06 - 3.55 | 308.42 | 10.95 – 668.06 | 64.98 |
| D10 | Moyar | 19.61 | 16.21 - 23.32 | 99.70 | 1.09 – 888.80 | 91.45 |
| D15 | Moyar | 6.09 | 5.79 - 6.40 | 55.57 | 0 – 910.11 | 194.62 |

The mean distance from home across all dogs was 74.44 ± 119.53 m (median = 36.12 m, range = 0 m - 1197.06 m) (Table 2). 91.8% of all GPS points (n=20453 of 22277) were within 250 m of the home, while 98.4% (n=21917) were within 500 m. The mean intensity of use was 115.94 ± 54.46 (range = 48.89 – 213.08, median = 91.45) (Table 2). When the three metrics, activity range, intensity of use, and mean distance travelled from home were compared with a Wilcoxon rank sum test with respect to sex, village, and day and night (the last used as a variable only for mean distance from home), the only significant difference was between intensity of use in each village, which was higher in Moyar than Masinagudi (Table 3, Fig. 3). The activity pattern analysis produced a relatively unimodal distribution, with a broad peak from the early morning hours (beginning at 1:00) until around noon with a dip at 6:00, followed by a steep decline until late night (Fig. 4). The collared dogs therefore displayed a partly diurnal, partly nocturnal activity pattern.

**Figure 3:**
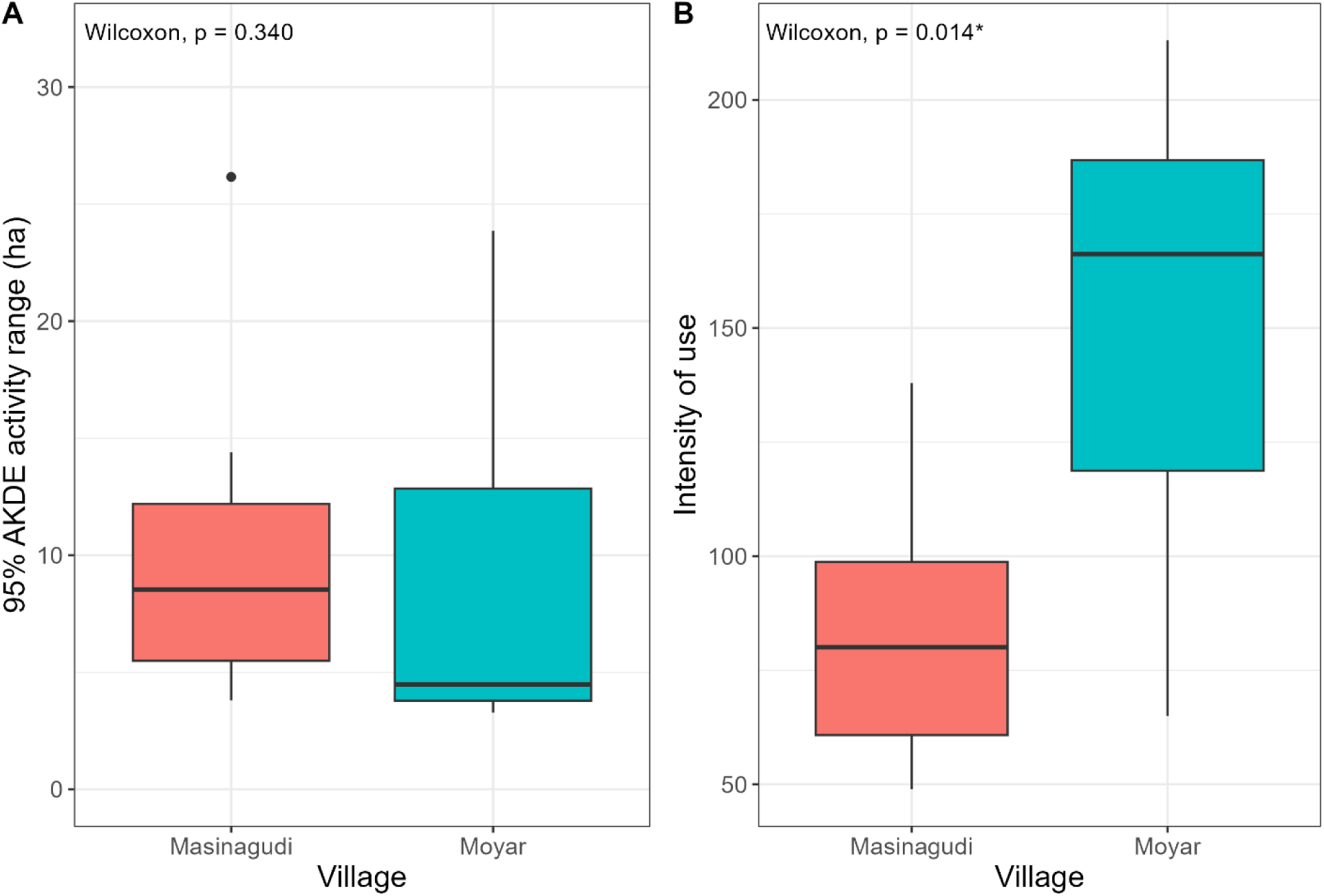
Box plots showing the differences in the home-range (A) and intensity of area use (B) for collared dogs from two villages (visualizing select values from Table 3). The values on the top left show the p-value from the Wilcoxon rank-sum test (significant values marked by an asterisk).

**Figure 4:**
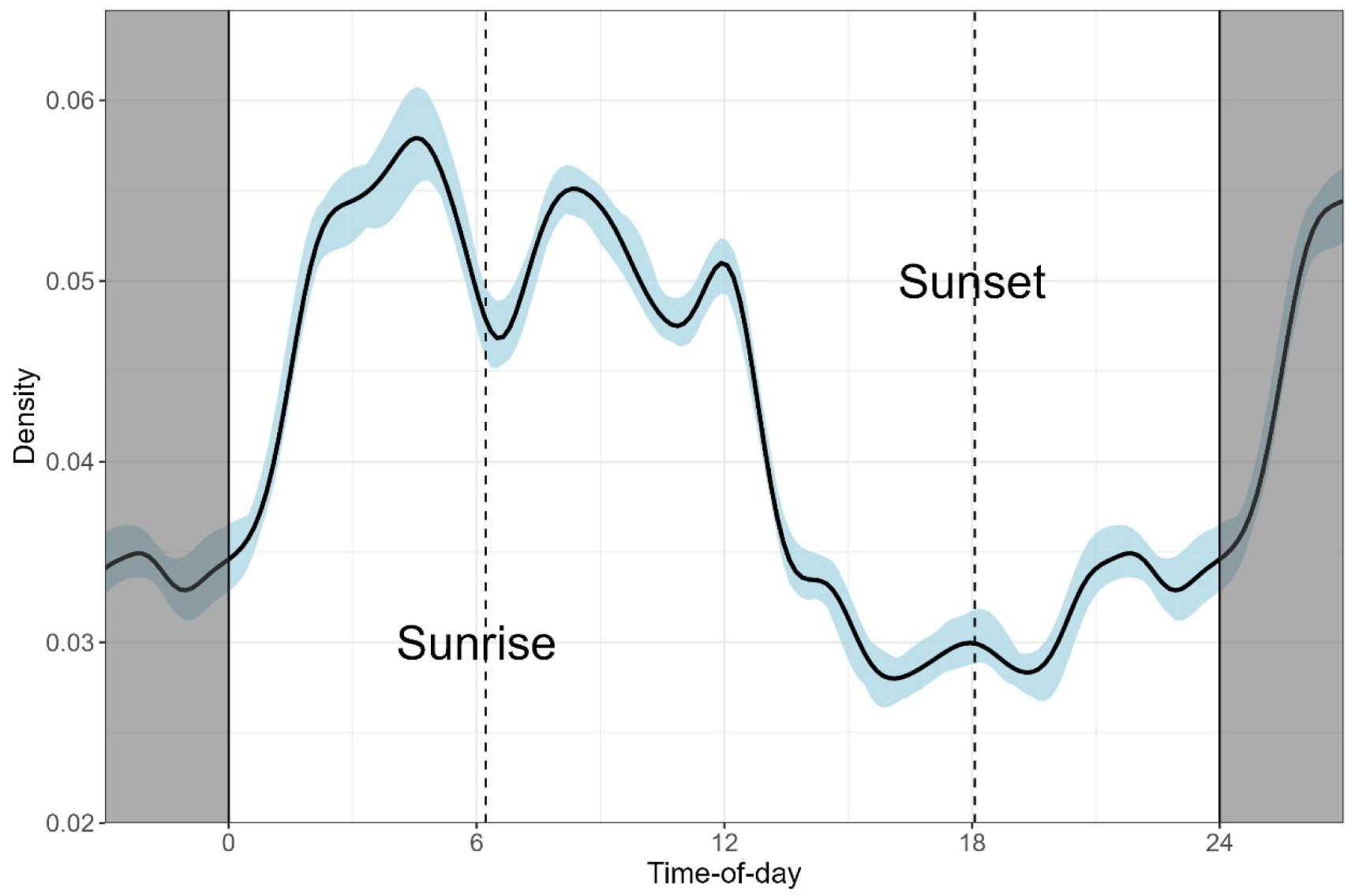
Kernel density plot showing collared FRD activity pattern across 24 hours.

**Table 3:** Wilcoxon test results for movement metric comparisons, with respect to different variables. Significant values (<0.05) are marked with an asterisk.

| Metric | Variable | P-value | U statistic |
| --- | --- | --- | --- |
| Mean distance to home vs<br>mean distance to centroid | NA | 0.486 | 130 |
| Mean distance to home | Day/night | 0.148 | 148 |
| Mean distance to home | Sex | 0.867 | 30 |
| Mean distance to home | Village | 0.397 | 20 |
| Activity range | Sex | 0.336 | 37 |
| Activity range | Village | 0.336 | 37 |
| Intensity of use | Sex | 0.121 | 42 |
| Intensity of use | Village | 0.014* | 7 |

The initial coarse assay of habitat selection found that 85.94% of all dog locations (n=19145) were in settlements, followed by 7.82% of locations (n=1742) in agricultural areas and 2.56% (n=571) in fallow/scrubland. Together, 96.32% of all locations were in human-settled or human-modified habitats. With 1.96% of locations (n=436) in deciduous forest and 1.23% (n=273) in thorn forest, 3.18% of all GPS locations were in the protected area. When examined by individual, 99.5% of the points located in the forest were from four FRD (Table S2, supplementary material), indicating that the other 11 either rarely ventured or did not venture into the forest at all during the sampling period. As the availability of settlements in the 70 sq. km landscape was 1.85%, while agriculture was 2.76% and fallow/scrub land was 2.56%, compared to 32.4% of deciduous forest and 59.89% of thorn forest, dogs strongly selected for human modified habitats over other land-use types.

This supported the results of the integrated step-selection analysis, which evaluated the fine-scale habitat selection of dogs from the study area. The outcome of iSSA was similar for both time scales (15 min, 30 min), with a higher standard error for the data resampled at 30 minutes (Fig. 5); therefore, we used the results of the 15 min resampled data for further analysis. Of all available habitat classes, anthropogenic habitats (that is, settlements and agricultural land) were significantly preferred, while others were not. However, the habitat-movement parameter interaction was significant for multiple habitats. The observed step-lengths were significantly longer than random steps in the forest habitat (β = 0.0092, p =0.043), whereas they were non-significant in agriculture (β = 0.0077, p =0.077) and fallow/scrub habitat (β = 0.0079, p =0.075). Overall, while the only significant habitat preference was for human-modified habitat classes, the movement speed in each class was substantially different, with FRD moving with the longest step lengths of approximately 85m-90m in the forested areas (traversing longer distances between GPS fixes), followed by fallow/scrubland and agricultural fields, where the step lengths were closer to 70m-75m (Fig. 6). The slowest movement with the shortest step length of 45-50m was recorded in settlements, and there was no significant difference when compared to random steps. Between villages, the movement trend was similar in both, but the movement speed was marginally higher in Moyar across all habitat classes (Fig. 6).

**Figure 5:**
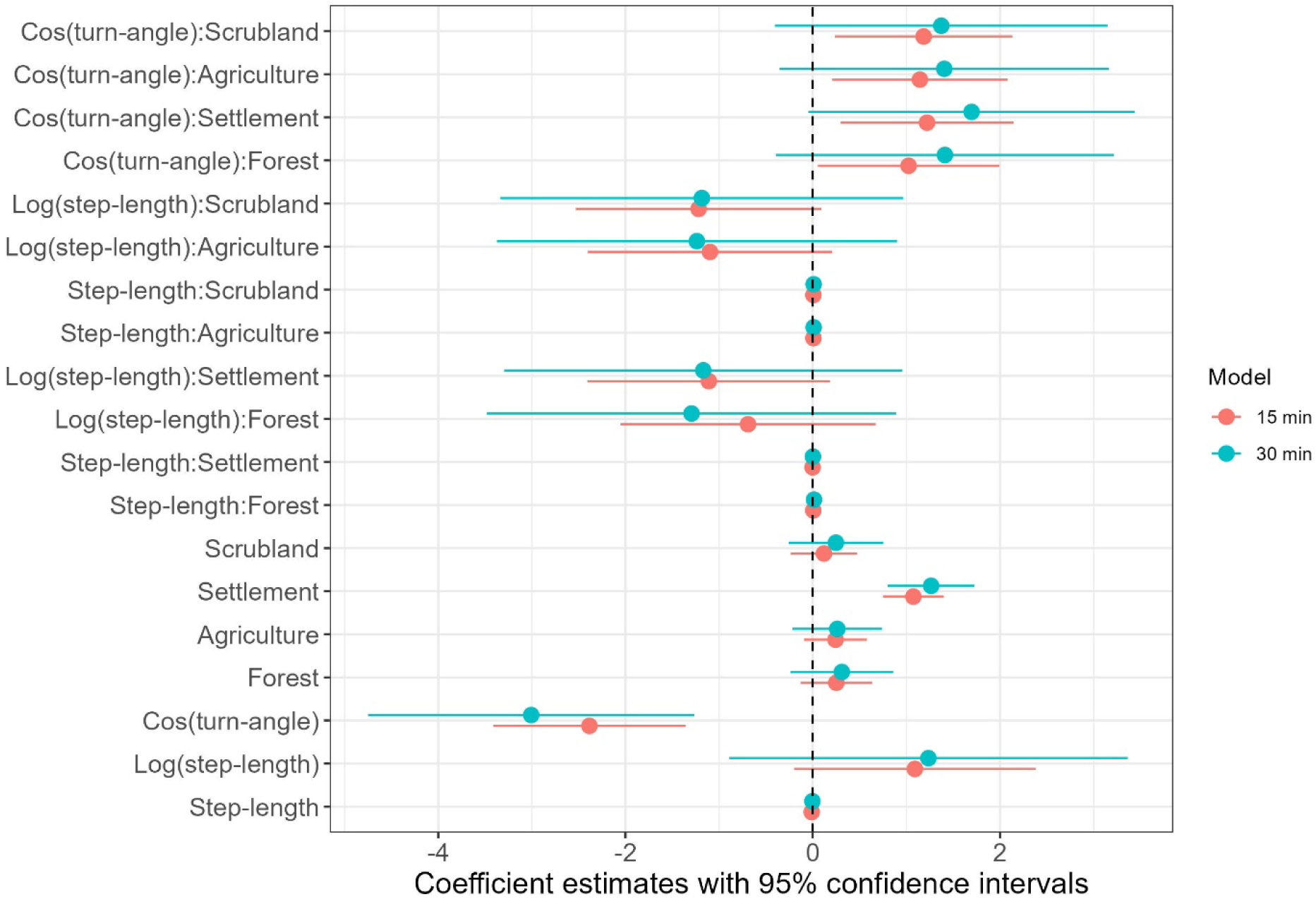
Coefficient estimates with 95% confidence interval from the integrated step-selection analysis for the two different temporal scales. The orange dots and lines represent the model where the temporal scale for consecutive fixes was resampled at 15 minutes, and the blue dots and lines represent the model where the temporal scale for consecutive fixes was resampled at 30 minutes.

**Figure 6:**
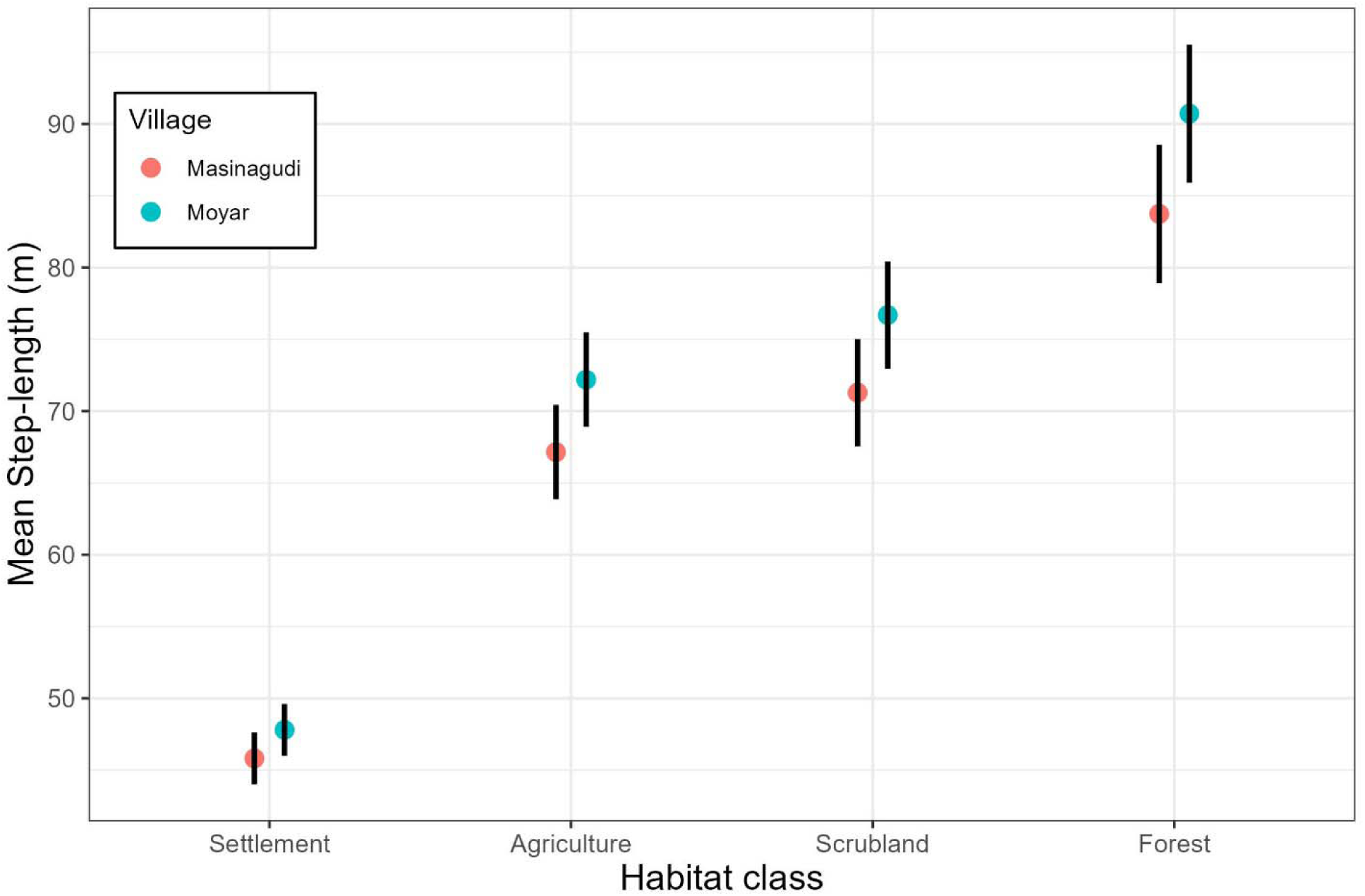
Estimates of mean step length (speed) for movement estimated for different habitats and different villages derived using the integrated step-selection function. The black bars indicate the 95% confidence interval for each of the habitat classes.

## Discussion

Our study investigates the movement ecology of owned free-ranging dogs in Mudumalai Tiger Reserve, India. We found that GPS-tracked FRD primarily selected for anthropogenic habitats, with an overwhelming preference for settlement areas, followed by agricultural land and fallow or scrubland at village boundaries. Overall, FRD utilized human-modified habitats more than 96% of the time and the area within 500 m of their owners’ homes more than 98% of the time.

This corroborates the well-documented phenomenon of dogs being closely associated with human-modified habitats in rural areas [31,52–54], and usually staying within a few hundred meters of their owner’s home [17,21,54]. Indeed, almost all dog movement in forested land could be attributed to only four FRD of the 15 that were tracked. Additionally, other movement metrics were low compared to estimates from other dog telemetry studies. For instance, the median activity range of 6.09 ha was much lower than median activity ranges from studies in Chad (54- 34 ha) [55], and Chile (19.20, 15.8-24.4) [17,56] (see also Table S2, supplementary material), while the maximum distance from home of 1197 m was fairly short compared to ranging distances recorded in Cambodia (up to 4km) [19] and Australia (8-30km) [14].

Given these limited movement patterns, the rural FRD in our study seem to fall into the class of *stay-at-home* dogs, according to the three classes defined by Hudson et al. (2017), unlike *roamers* or *explorers*, both of which travel to areas distant from owners’ home with notable frequency. As the FRD in MTR displayed minimal roaming and consistently used the area immediately surrounding their owners’ homes, they are likely to have the majority of their requirements fulfilled by the humans they depend on [13]. This was an expected finding, since we collared owned FRD which were subsidized by their owners, and indicates that this class of dogs is unlikely to pose a persistent threat to wildlife. However, although our results are relevant to local dog management in MTR, as the vast majority of the dog population in villages on the Sigur Plateau are owned and sterilized (SVK, unpub. data), we note that we have not investigated the movement of feral or non-owned dogs, and cannot comment on their potential ranging into forest areas or impact on wildlife.

Our study was also constrained by a few other limitations, mainly owner apprehensions resulting in the unauthorized removal and subsequent loss of collars, and collar malfunction resulting in data loss, which reduced our effective sample size from 26 dogs to 15 dogs and limited the sampling period. While several telemetry studies have been conducted on dogs with shorter sampling periods (see Table S2, supplementary material), we acknowledge that the sampling period should ideally be longer to more thoroughly investigate dog behavior and distinguish between short-term and long-term movement patterns; for instance, dog home ranges have been found to vary significantly across seasons [55]. Additionally, since we were unaware of when dogs accompanied humans, we could not differentiate between natural movement versus movement that reflected human activities (such as working in agricultural fields). However, we believe this is unlikely to have significantly impacted our results, as many owners reported that their dogs did not always accompany them but roamed freely on their own. This is supported by the kernel density activity pattern, which showed that dogs were generally more active than humans between 0:00-6:00 and less active than humans between 15:00-19:00.

This activity pattern also indicates why we found no significant difference between day and night mean distance from home. While we expected that dogs would move less at night to minimize chances of encountering wildlife, as predator species are more active at village boundaries after sunset, the activity pattern showed that FRD moved during daylight as well as nighttime, with a peak between early morning hours before dawn until around noon. This pattern differs from the bimodal (morning-evening) activity pattern displayed commonly displayed by dogs elsewhere [57] but tallies with the dog activity pattern in areas with puma presence [58]. If the observed pattern is due to the close proximity to wildlife-adjacent areas, FRD may have adapted to predators in MTR being more active late in the night and decreasing their activity closer to dawn, or they may may derive sufficient protection from living in settled areas to be active at night; however, confirming either theory would require data on wild predator activity patterns. We also found no difference in movement metrics with respect to sex, unlike in other studies, where neutered males have been recorded as having significantly larger activity ranges than spayed females [45]; this may also be due to the presence of predators, as they may constrain male FRD to similar activity ranges and distance travelled from home as female FRD.

In contrast, the significant difference in intensity of use between villages indicates that FRD used their activity ranges differently in each village. In Moyar, the human population is smaller and village homes are divided into settlement patches separated by large stretches of agricultural land, while the human population in Masinagudi is several times larger and agricultural patches are fragmented by well-connected roads and densely clustered houses. Therefore, we interpret the higher intensity of use by Moyar FRD, indicating that they travelled longer distances or more frequently within activity ranges similar to those of Masinagudi FRD, as dogs having to move between safe, well-known areas inside their activity ranges in search of refuge from wildlife encounters. Searching for resources or mates is unlikely to be a significant factor here as dogs were fed at home by their owners and almost all dogs were sterilized. The higher intensity of use by Moyar FRD is likely due to the heightened predation risk posed by wildlife, stemming from two factors: the smaller human population with fragmented clusters of settlements, and the larger, more porous perimeter with the forest and swathes of open agricultural land inside the village that are conducive to wild animal movement. Indeed, several wild species, including leopards, have previously been camera trapped in such land inside Moyar (SVK, unpub. data). In Masinagudi, on the other hand, the larger human population, smaller village-forest boundary and increased anthropogenic infrastructure potentially provide more relative safety to FRD throughout their activity ranges, allowing FRD to carry out their daily activities with less total movement. That the FRD of Moyar and Masinagudi adopted different roaming behaviors, even when the two study sites were situated within 5km of each other, indicates that FRD can display a high degree of behavioral plasticity depending on fine-scale ecological context.

The results of the integrated step selection analysis, namely the significant difference in step length between in different habitats, also support the hypothesis that FRD rely on the human shield and perceive higher risk in forest land. Forest areas were traversed almost two times faster than settlements, and about 1.2 times faster agricultural or scrubland areas. This indicates that forest areas were used in a more utilitarian manner (perhaps for travel between points) while settlements were used for rest or play. Movement in agricultural and scrubland areas may have been exploratory forays or may have been while accompanying owners. Additionally, the higher movement speed across all habitat classes in the less populated and more fragmented village of Moyar, while not statistically significant, corroborates our interpretation of the difference in intensity of use between villages as being due to the difference in potential for wildlife encounters. It also suggests that average dog movement speed could potentially be inversely proportional to the settlement level in areas where FRD are preyed on by wildlife; that is, dogs may move slower in more built-up areas and faster in less built-up areas in response the level of perceived threat, although confirming this would require studying FRD across more villages with a larger range of settlement levels.

Overall, we recommend that studies documenting patterns of dog movement should be undertaken before designing and implementing dog management strategies that aim to modify roaming behavior, as they may uncover similar unexpected nuances based on human-mediated factors. For such studies, we suggest collaring more broadly across dog demographics (namely ownership, sex, age and sterilization status) as well as across human and village related parameters (e.g., length of the village-forest perimeter, household density, agricultural cover) at larger spatial and temporal scales. In particular, studies from non-protected areas that support wildlife may not only be easier to carry out but may provide much-needed insight into the dog-wildlife interface in areas with no restrictions on human movement and land use.

## Conclusion

Using movement metrics and the robust framework of integrated step-selection analysis to model dog habitat selection, we compared free-ranging dog movement between two sites at a fine geographical scale, interpreted through the lens of the human shield hypothesis [49]. Our findings, while not generalizable across all dog-wildlife interfaces and limited by a relatively short sampling period, contribute to research on dog movement and space use by documenting metrics such as the activity range, and showing that FRD display high behavioral plasticity, as movement patterns can differ between adjacent villages based on the level of protection afforded by human settlements, putatively to avoid wild predators. They also show that owned, provisioned and sterilized free-ranging dogs from villages in MTR spend the vast majority of their time inside human settlement areas, making their potential impact on wildlife relatively low compared to other dog classes, such as feral dogs. Differential access to beneficial resources – in this case, refuge from wildlife encounters — provided by the intensity of human land use should be taken into account when managing dog movement and behavior in wildlife adjacent areas, given their highly commensal relationship with humans.

## Supporting information

Revised additional file 1

New additional file 2

## Declarations

### Ethics approval and consent to participate

All GPS collars were fitted with the owners’ consent. Collars weighed less than 250 g and were within the required weight limit of < 5% of the dog’s bodyweight, causing no physical distress to dogs during the duration of deployment.

### Consent for publication

Not applicable

### Availability of data and materials

The dataset generated during this study is available in the Movebank repository under the study name “Free-ranging dogs - vksanj - Mudumalai TR - GPS collaring”, study ID 5328251712, at the following link - https://www.movebank.org/cms/webapp?gwt_fragment=page=studies,path=study5328251712

Codes for data analysis are available on Github at the following link - https://github.com/nilanjanchatterjee/Dog-population-and-movement-Analysis/blob/main/02_Movement_analysis.html

### Competing interests

The authors declare that they have no competing interests.

### Funding

This study was funded by The Habitats Trust (Seed Grant, March 2024) in a grant provided to SVK.

### Authors’ contributions

SVK acquired funding for the study, conducted fieldwork, participated in data analysis, and drafted the manuscript. NC led data analysis and contributed to manuscript writing. RK provided logistic and supervisory support, and contributed to manuscript writing. All authors read and approved the final manuscript.

## Acknowledgements

We acknowledge and appreciate the support provided by staff at the Indian Institute of Science field station in Masinagudi, namely Mr. Sharan B. Gowda, Mr. Albert Suresh, Mr. Bharanaiah, Mr. Arjunan, and Dr. Suresh H. S. We are grateful to Mr. Arasu for field support, as well as the dog owners of Masinagudi and Moyar for permission to collar their dogs. We thank two anonymous reviewers for their comments that helped improve this manuscript.

## Additional files

Additional file 1, format: PDF Data:

Table S1: Number of GPS points for each individual dog in each habitat type.

Table S2: Home/activity range comparisons from global studies — expanded table adopted from Ladd et al., 2023, with home/activity range estimates and sampling durations from dog telemetry studies across countries.

Table S3: ΔAIC comparison across activity range models — ΔAIC values for all trialed models during activity range estimation.

Figure S1. Time plots for GPS locations in the forest, illustrating individual temporal variation in dogs’ use of forest land.

Figure S2. Compilation of photos of collared dogs.

Additional file 2: Mapped kernel density estimates for all dogs. Please note that this includes data from all 22 dogs uploaded to Movebank, and dogs are referred to by names and not ID numbers.

## Notes

https://www.movebank.org/cms/webapp?gwt_fragment=page%3Dstudies%2Cpath%3Dstudy5328251712443

https://github.com/nilanjanchatterjee/Dog-population-and-movement-Analysis/blob/main/02_Movement_analysis.html

### Competing Interest Statement

The authors have declared no competing interest.

### Summary of Updates

Author affiliation updated, supplementary material expanded, Figures 1 and 2 revised, discussion and conclusion substantially revised to limit the inferences drawn, abstract, introduction, methods and results revised to a minor extent to better reflect the altered focus of the paper.

