## Supplementary material for "Of Shared Homes and Pathways: Free-Ranging Dog Movement and Habitat Use in a Human-Wildlife Landscape in India": Revised additional file 1

Table S1. Number of GPS points for each individual dog in each habitat type. Dog names are provided to facilitate reference to the Movebank database.

| ID | Name | Agricultural  land | Deciduous forest | Fallow/  scrubland | Settlement | Thorn forest | Water | Total |
| --- | --- | --- | --- | --- | --- | --- | --- | --- |
| D1 | Appu | 22 |  | 45 | 1102 |  |  | 1169 |
| D2 | Biscuit | 36 |  | 27 | 2365 |  | 18 | 2446 |
| D3 | Brown | 561 |  | 45 | 2176 | 1 |  | 2783 |
| D4 | Burger | 301 | 202 | 10 | 1638 | 21 |  | 2172 |
| D5 | Chan | 490 | 30 |  | 1303 |  | 51 | 1874 |
| D6 | Kamaru | 36 |  | 113 | 731 | 25 | 1 | 906 |
| D7 | Kutti | 9 | 1 | 3 | 736 | 1 | 7 | 757 |
| D8 | Mani | 44 |  | 3 | 2127 |  |  | 2174 |
| D9 | Meenu | 1 |  | 3 | 643 |  |  | 647 |
| D10 | Puppy | 134 | 203 | 26 | 622 | 219 | 3 | 1207 |
| D11 | PuppyMS | 35 |  |  | 1105 |  |  | 1140 |
| D12 | Simba | 8 |  | 64 | 1428 | 1 |  | 1501 |
| D13 | Sophie | 19 |  | 19 | 581 |  |  | 619 |
| D14 | Sundari | 28 |  | 5 | 602 |  |  | 635 |
| D15 | Vellian | 18 |  | 208 | 1986 | 5 | 30 | 2247 |
|  |  | 1742 | 436 | 571 | 19145 | 273 | 110 | 22277 |

Table S2. Table of median home/activity range from studies across different countries (adapted from Ladd et al., 2023; rows marked with an asterisk are new additions to the table.)

| Study | Location | Method | Median home/activity range (ha) | Tracking  duration | Tracking  method |
| --- | --- | --- | --- | --- | --- |
| Meek (1999) | Australia | MCP isopleth, outliers excluded | 72.50 | 15 months | VHF radio tracking |
| Van Kesteren et al. (2013) | Kyrgyzstan | Characteristic hull polygon | 2.26 | Mean 20 h | GPS tracking |
| Sparkes et al. (2014) | Australia | MCP 100% isopleth, forays excluded | 37.47 | 7 days | GPS tracking |
| Dürr and Ward (2014) | Australia | MCP 95% isopleth | 3.60 | Mean 50 h | GPS tracking |
| Ruiz-Izaguirre et al. (2015) | Mexico | Kernel density | 16.10 | 45 days | VHF radio tracking |
| Kennedy et al. (2018) | Australia | MCP 100% isopleth, forays excluded | 8.88 | 3–5 days | GPS tracking |
| Muinde et al. (2021) | Kenya | MCP 95% isopleth | 9.30 | 5 days | GPS tracking |
| Saavedra-Aracena et al. (2021) | Chile | Kernel density | 19.20 | Mean 20.5 days | GPS tracking |
| Warembourg et al. (2021) | Chad  Guatemala  Indonesia  Uganda | Biased random bridge, 95% isopleth | 7.70  5.70  5.60  5.70 | Median 60.3 h | GPS tracking |
| Ladd et al. (2023) | Cambodia | MCP 95% isopleth | 91.40 | Mean 25.5 days | GPS tracking |
| Wilson-Aggarwal et al. (2021)* | Chad | Continuous time movement models, AKDE (95% | 54 (dry season)  31 (wet season) | Mean 37 days | GPS tracking |
| Dürr et al. (2017)* | Australia | Biased random bridge, 95% isopleth | 4.48 | 2-16 days | GPS tracking |
| Hudson et al. (2017)* | Australia | Biased random bridge, 95% isopleth | 5.59 | Median 13 days | GPS tracking |
| Molloy et al. (2017)* | Australia | Biased random bridge, 95% isopleth | 3.1 | Median 25 hours | GPS tracking |
| Schuttler et al. (2022)* | Chile | Continuous time movement models, AKDE (95% | 15.8 (spring)  21 (summer)  24.4 (autumn)  16.2 (winter) | Median 19 days | GPS tracking |
| Current study* | India | Continuous time movement models, AKDE (95% | 6.09 | Median 9 days | GPS tracking |

Table. S3. ΔAIC values for the activity range model comparisons for each individual dog.

| Dog ID | OU anisotropic | OUF anisotropic | OU | OUF | OUf anisotropic | IID anisotropic |
| --- | --- | --- | --- | --- | --- | --- |
| D1V1 | 0 | 2.009957 | 13.75997 | 15.76301 | 420.035957 | 2246.50817 |
| D2V2 | 0 | 2.006023 | 21.6967 | 23.6997 | 342.939535 | 1497.724371 |
| D3V2 | 0 | 2.001582 | 247.3654 | 249.3635 | 758.298947 | 3649.126524 |
| D4V2 | 0 | 1.998544 | 20.78438 | 22.77921 | 1211.907196 | 5141.972932 |
| D5V2 | 0 | 2.009705 | 32.2696 | 34.27499 | 58.811705 | 392.96917 |
| D6V1 | 0 | 2.004244 | 7.781406 | 9.776983 | 677.985195 | 2849.912699 |
| D7V2 | 0 | 2.022672 | 25.91456 | 111.8829 | 113.893971 | 203.612247 |
| D8V1 | 0 | 2.303314 | 96.33015 | 104.3857 | 177.875656 | 1122.994657 |
| D9V1 | 0 | 1.992732 | 21.40403 | 23.392 | 197.887928 | 351.337341 |
| D10V2 | 0 | 2.422494 | 11.09954 | 13.76486 | 977.961256 | 4103.394207 |
| D11V1 | 0 | 2.016833 | 25.11982 | 108.9501 | 110.959721 | 141.017389 |
| D12V1 | 0 | 2.004023 | 15.12266 | 17.12196 | 519.492014 | 1845.546518 |
| D13V1 | 0 | 1.965725 | 10.22088 | 12.17643 | 587.676573 | 1155.341411 |
| D14V1 | 0 | 2.010527 | 34.93571 | 36.9367 | 213.107512 | 646.151481 |
| D15V2 | 0 | 2.007764 | 128.2206 | 130.2247 | 133.282096 | 1017.879314 |

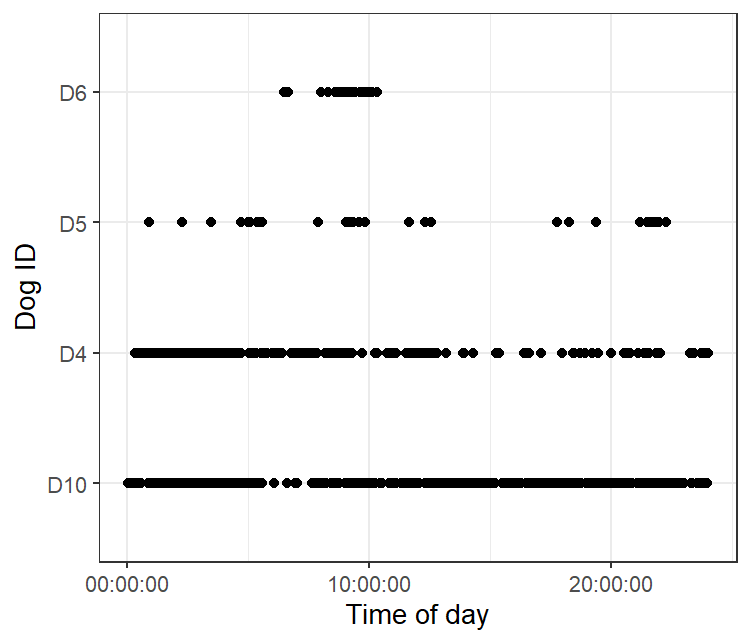

Figure S1. Time plots for GPS locations in the forest, illustrating individual temporal variation in dogs’ use of forest land. Here, D6 is a female sterilized shepherd’s dog, and D4 is a male unsterilized dog. D5 and D10 are female and male respectively, both sterilized. No other dogs utilised forest land in this study.

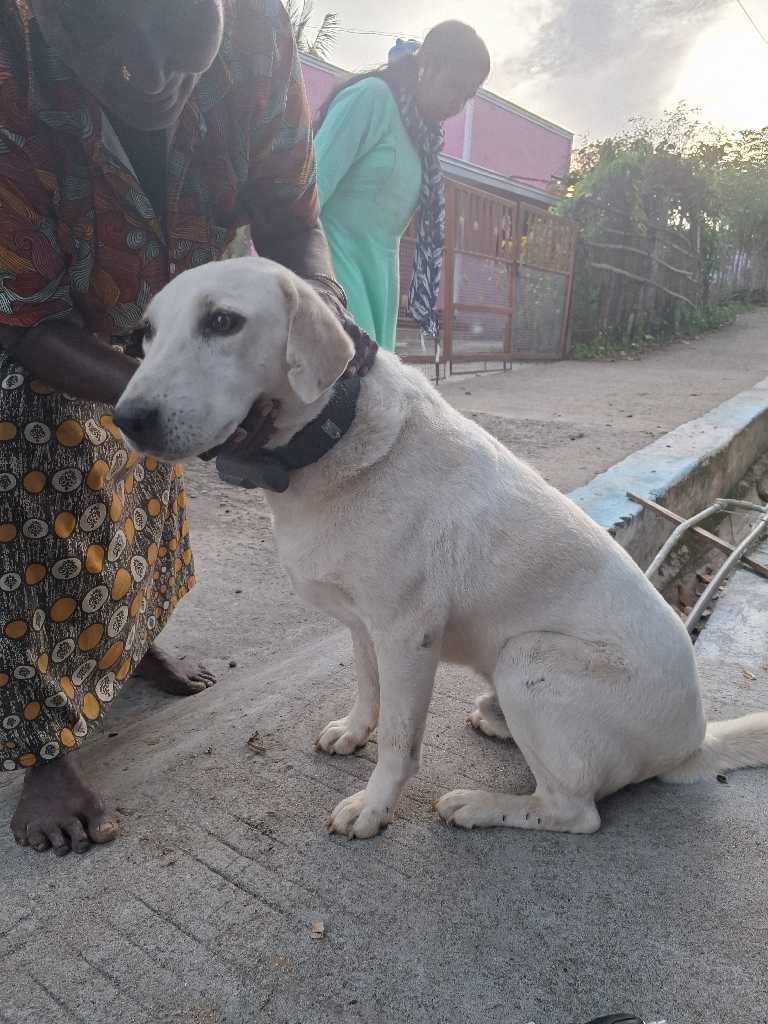

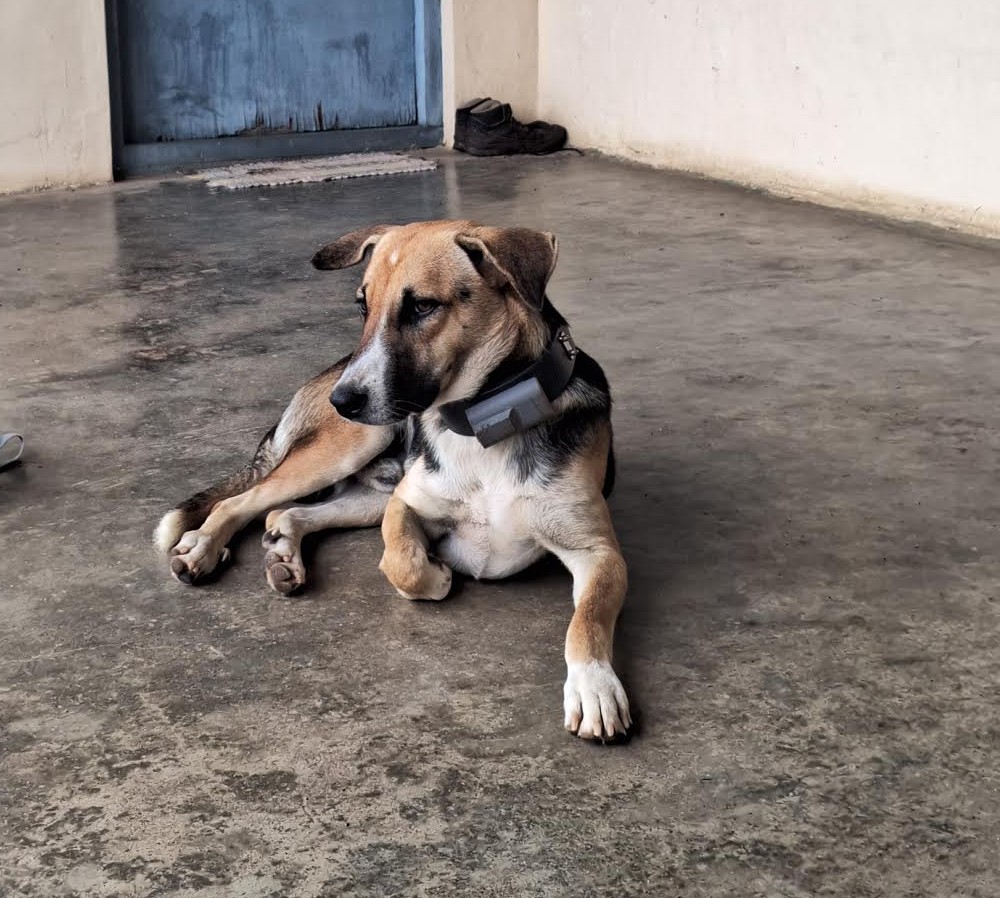

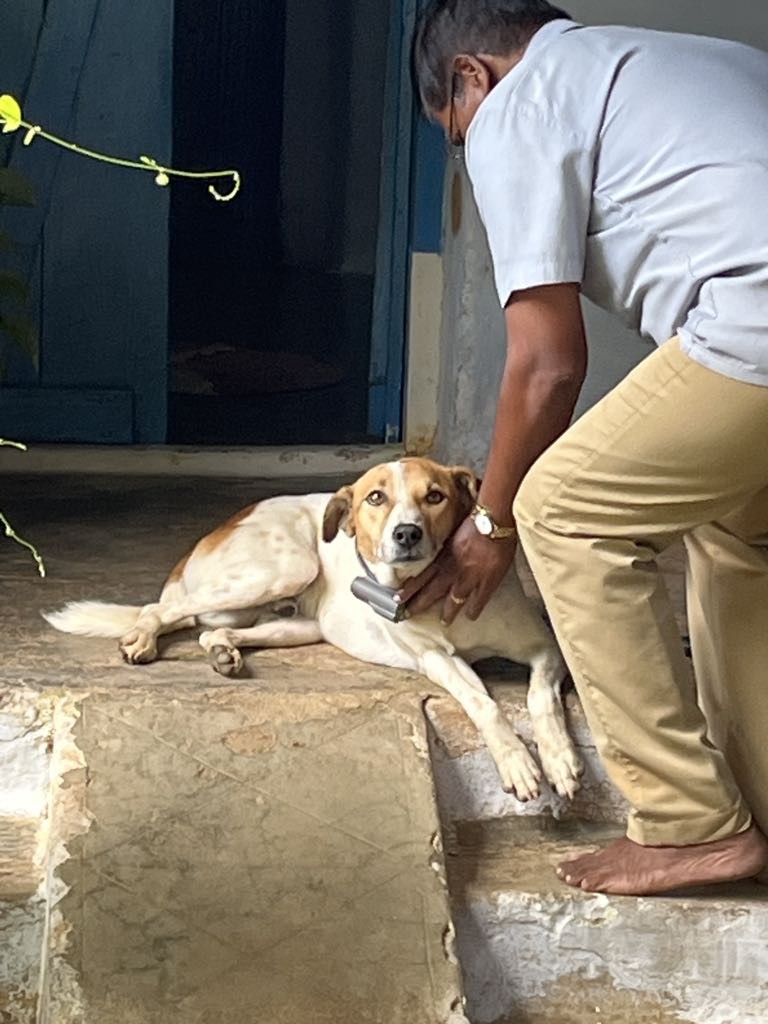

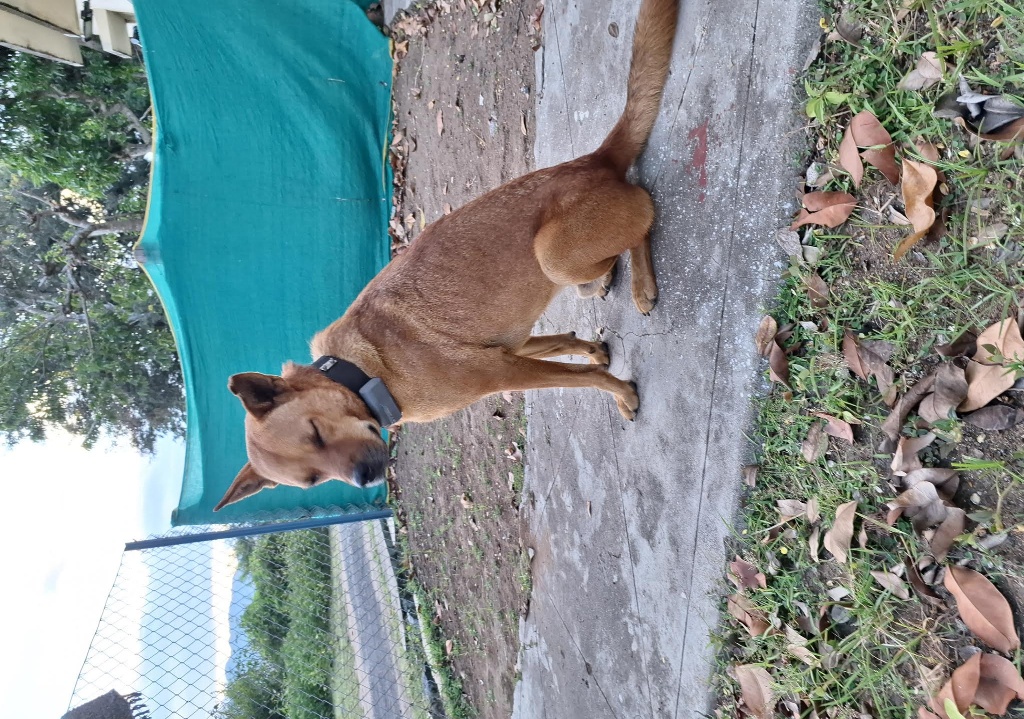

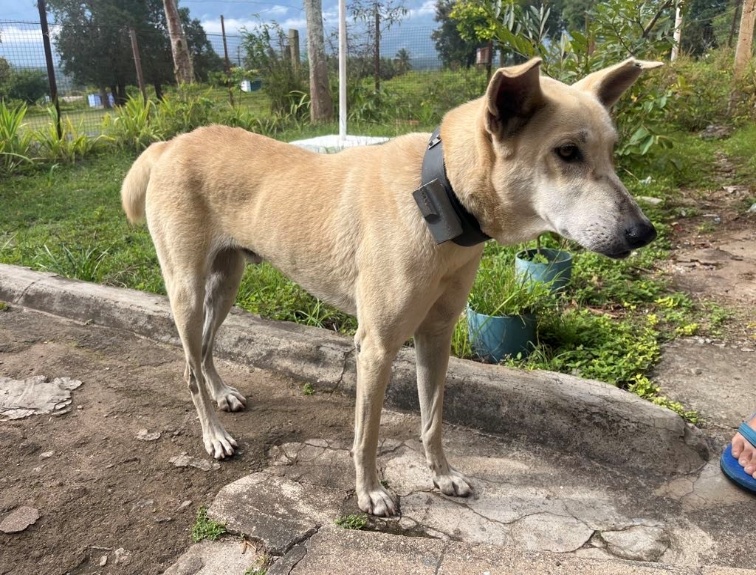

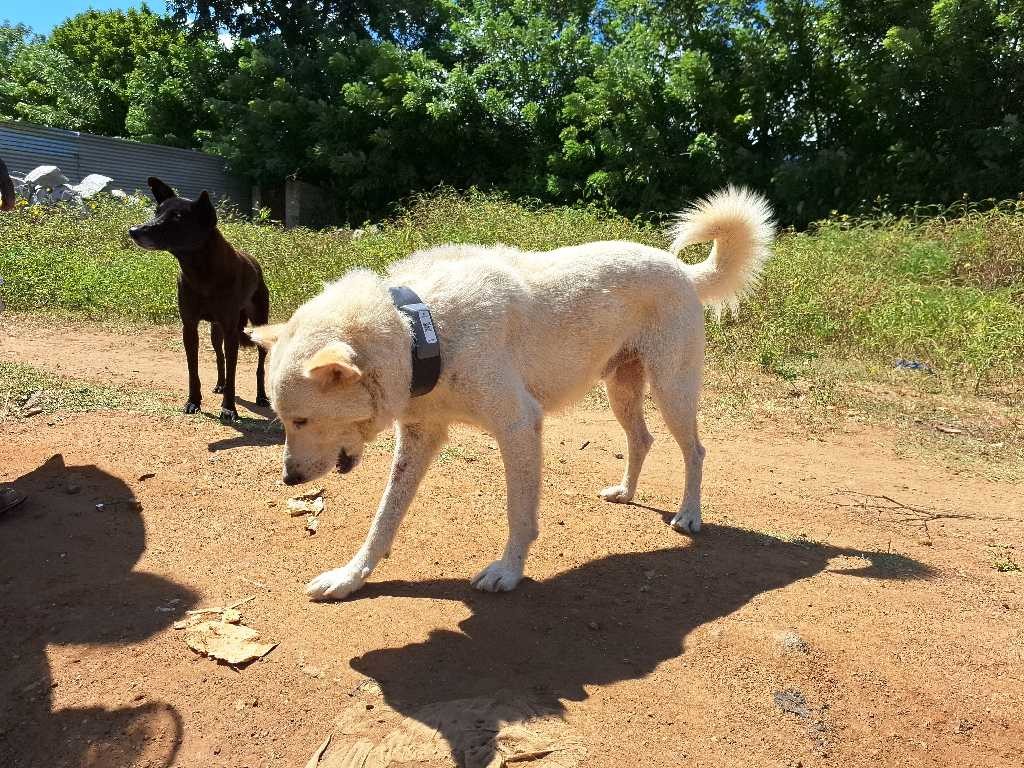

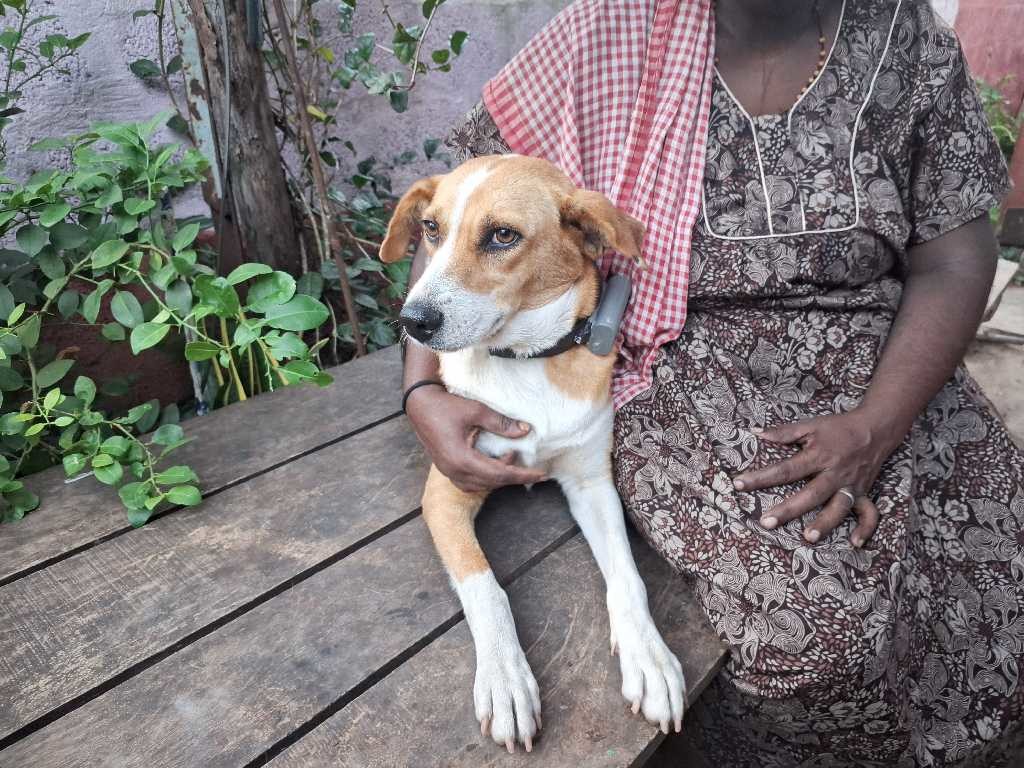

F

E

D

C

B

A

Fig S2. Photographs of collared dogs (A-F).
