## Supplementary material for "Of Shared Homes and Pathways: Free-Ranging Dog Movement and Habitat Use in a Human-Wildlife Landscape in India": New additional file 2

### Mani

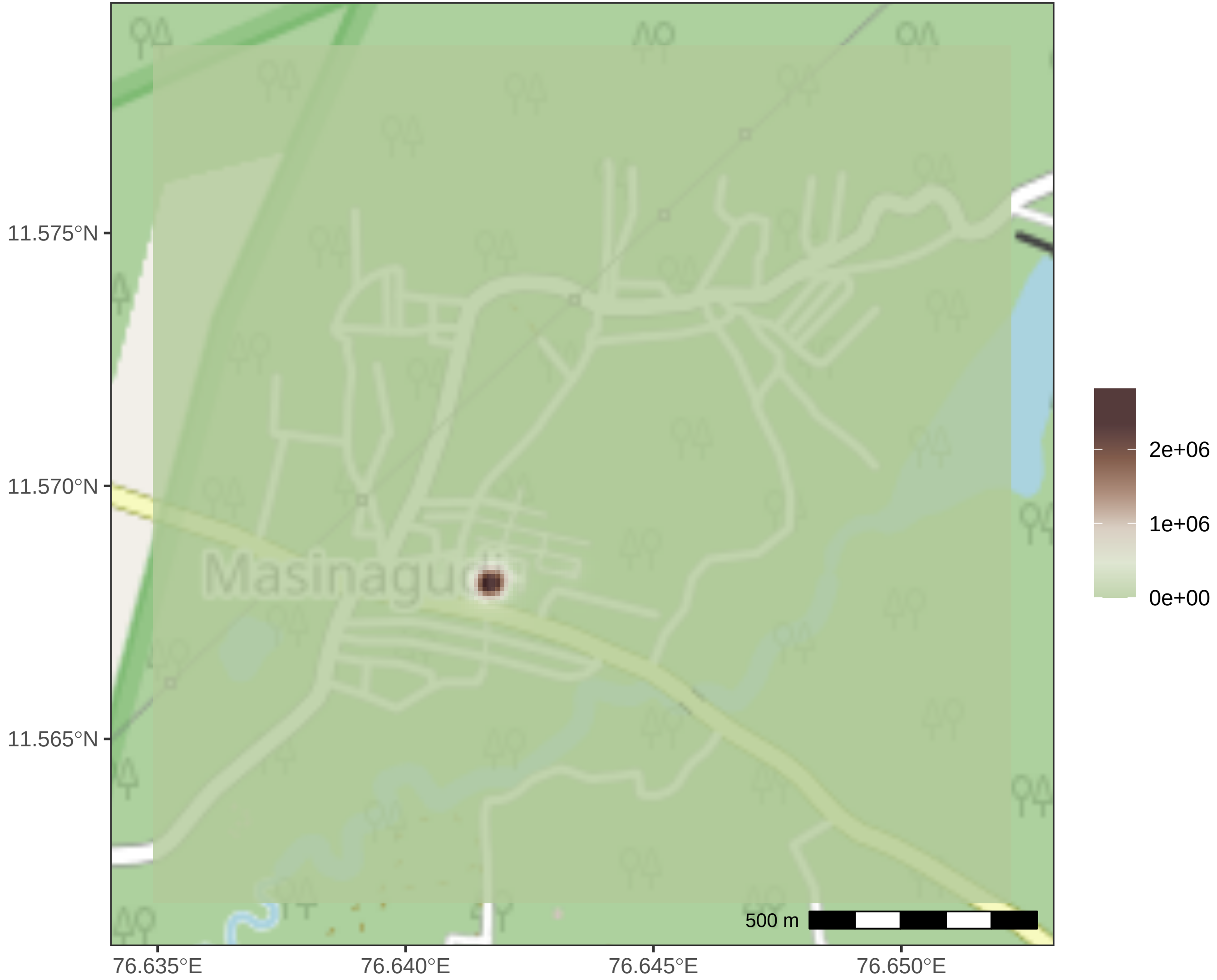

Masinagudi
500 m
11.575°N
11.570°N
11.565°N
76.635°E
76.640°E
76.645°E
76.650°E
2e+06
1e+06
0e+00

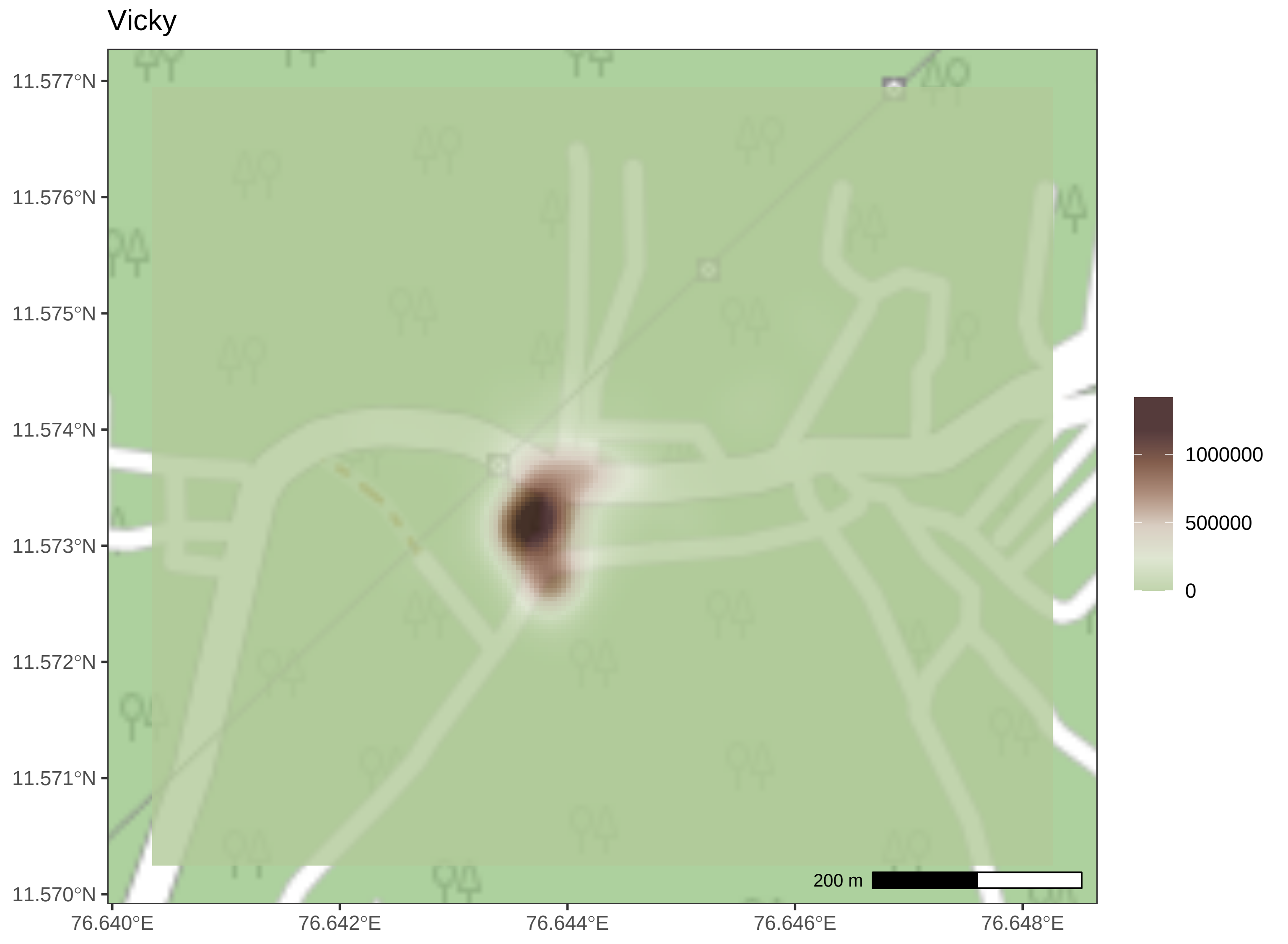
Vicky
11.577°N
11.576°N
11.575°N
11.574°N
11.573°N
11.572°N
11.571°N
11.570°N
76.640°E
76.642°E
76.644°E
76.646°E
76.648°E
200 m
1000000
500000
0

Sundari

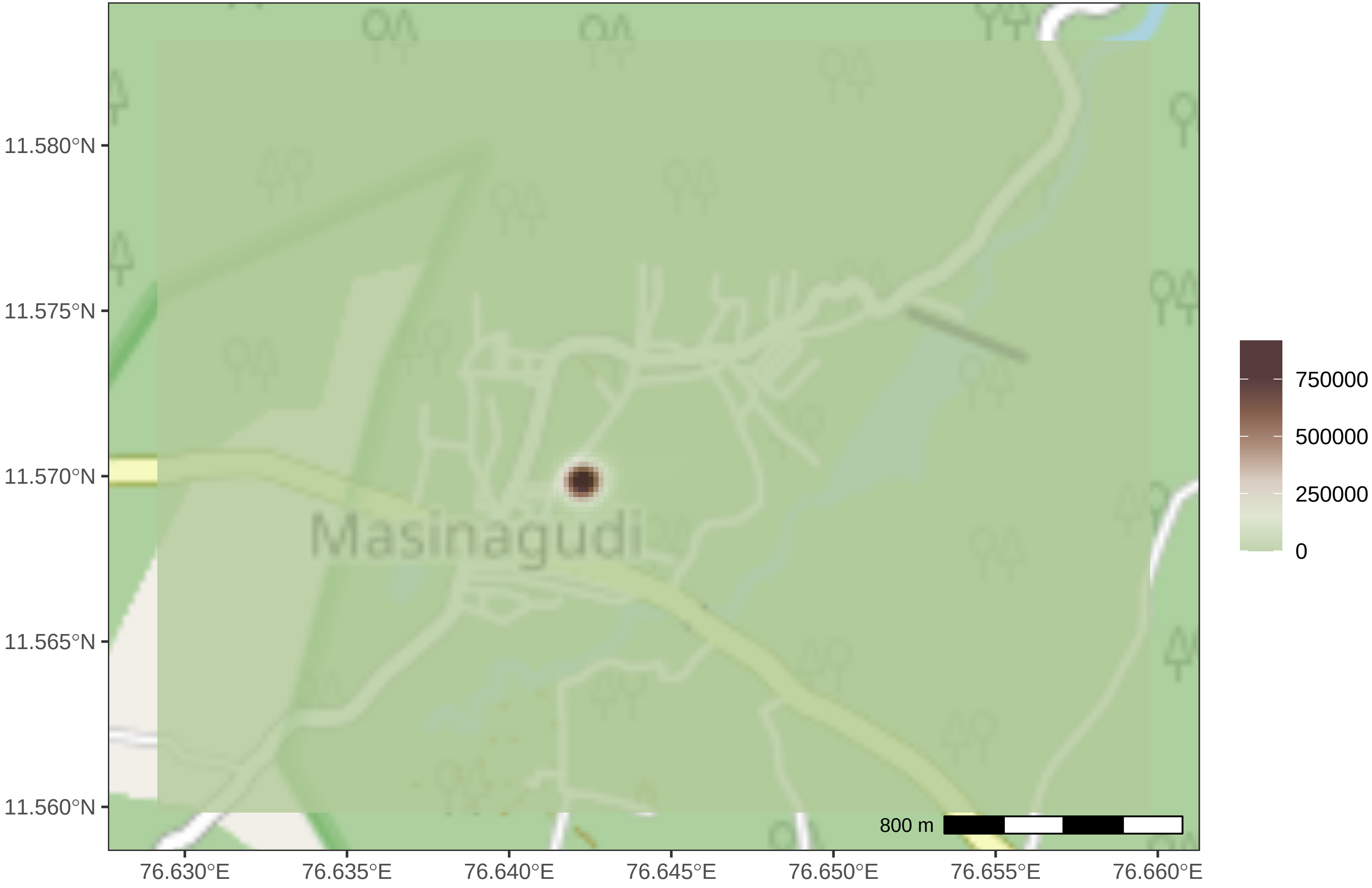

11.580°N
11.575°N
11.570°N
11.565°N
11.560°N
76.630°E
76.635°E
76.640°E
76.645°E
76.650°E
76.655°E
76.660°E
Masinagudi
800 m
750000
500000
250000
0

Pied

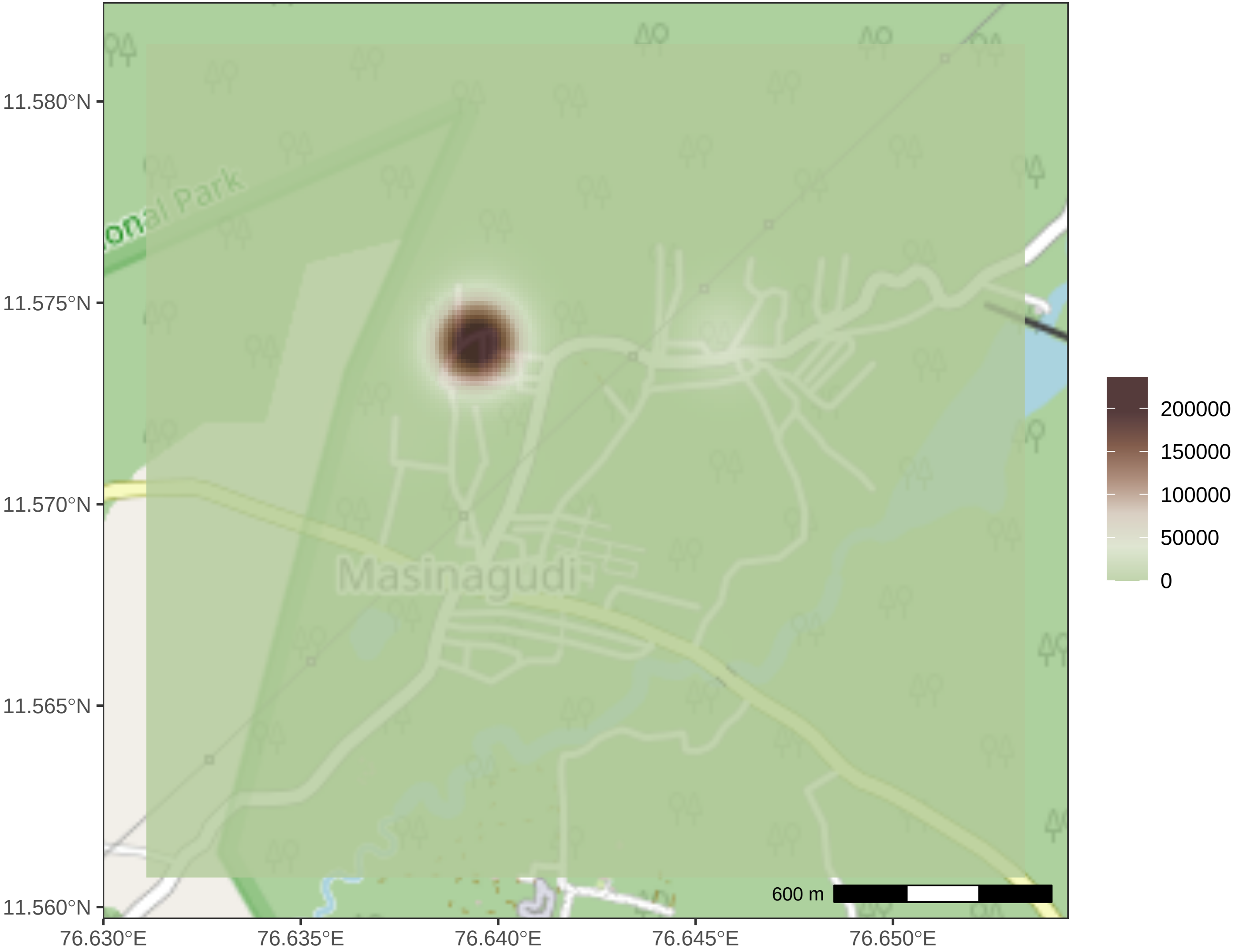

11.580°N
11.575°N
11.570°N
11.565°N
11.560°N
76.630°E
76.635°E
76.640°E
76.645°E
76.650°E
Masinagudi
onal Park
600 m
200000
150000
100000
50000
0

Kamaru

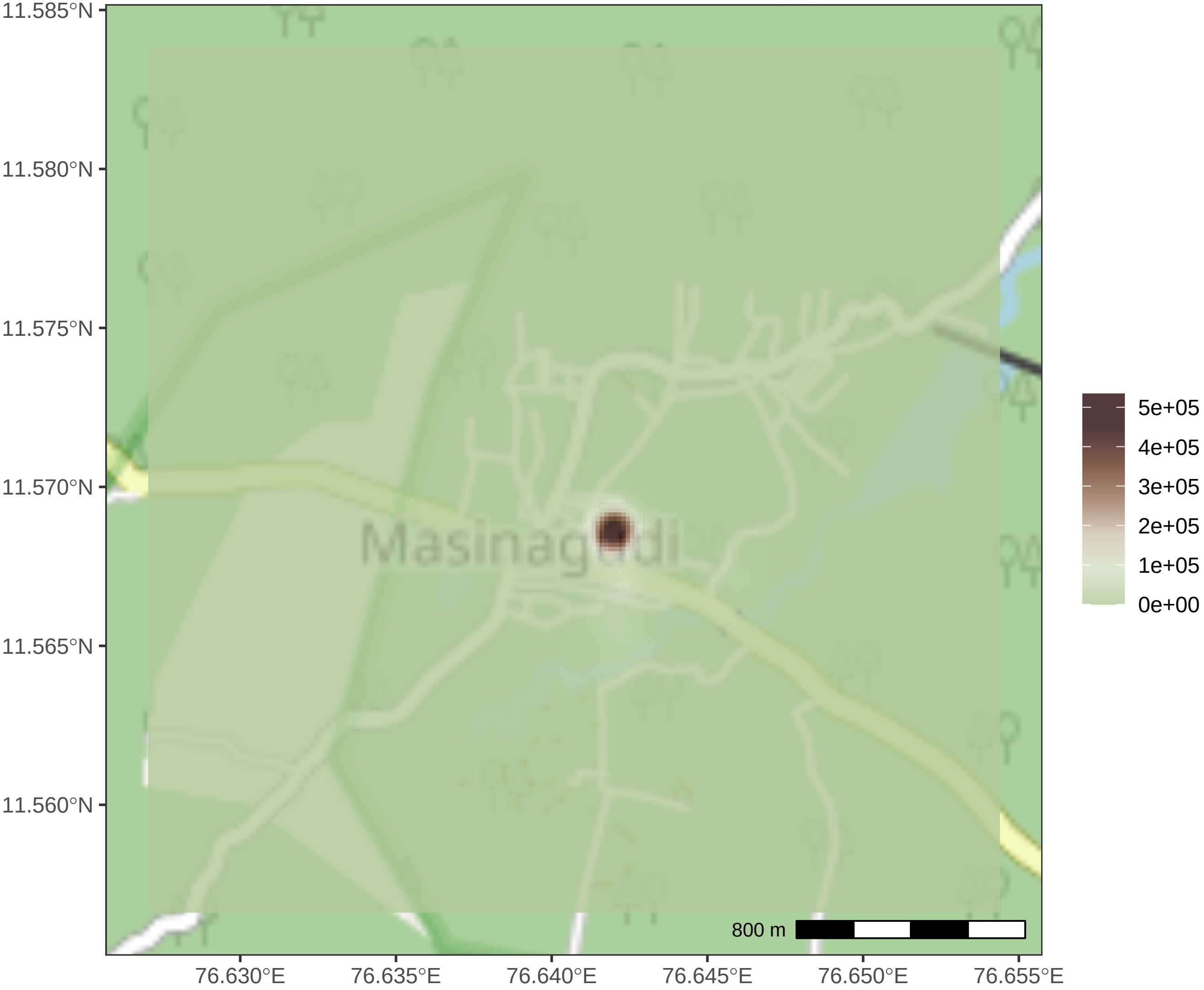

11.585°N
11.580°N
11.575°N
11.570°N
11.565°N
11.560°N
76.630°E
76.635°E
76.640°E
76.645°E
76.650°E
76.655°E
Masinagudi
800 m
5e+05
4e+05
3e+05
2e+05
1e+05
0e+00

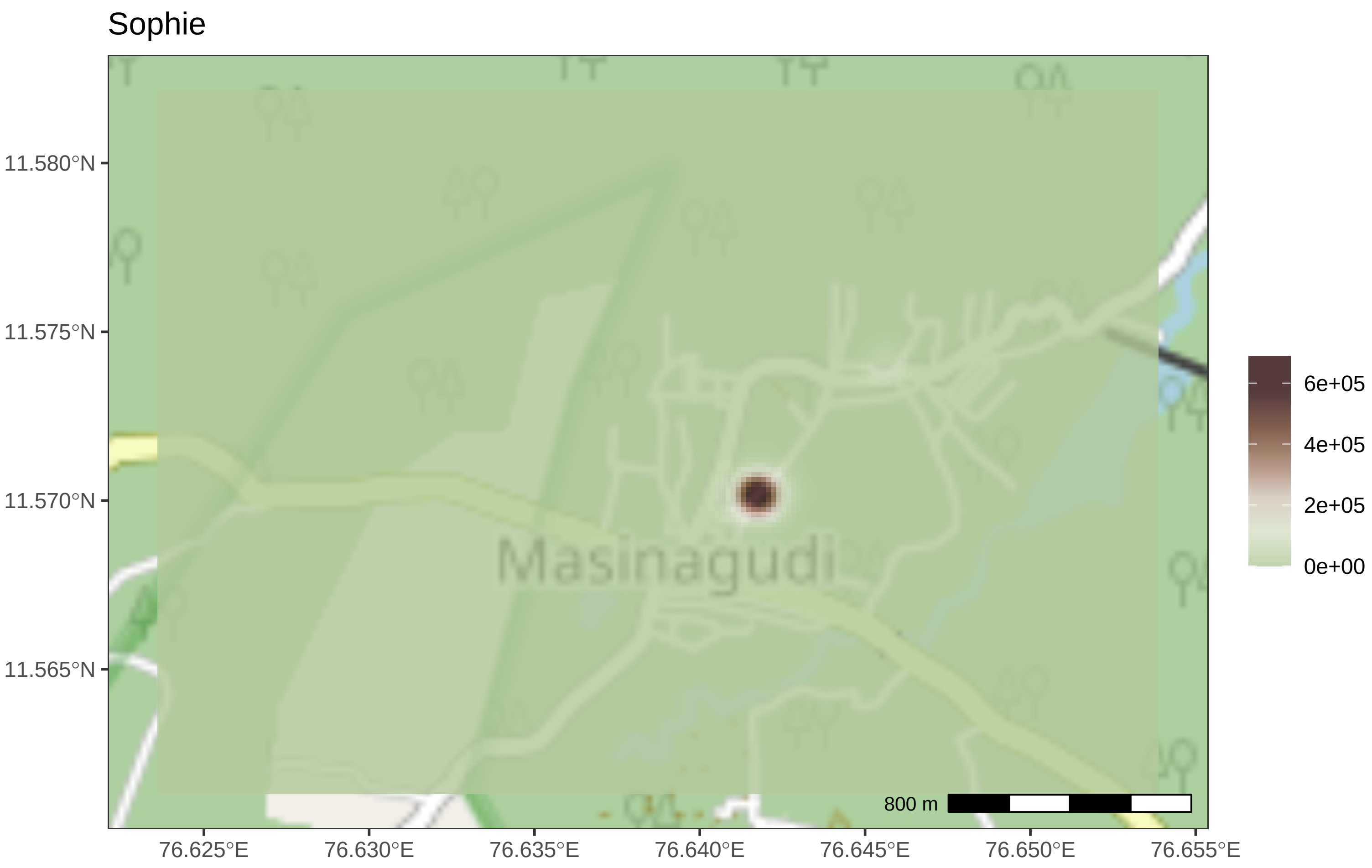

Sophie
11.580°N
11.575°N
11.570°N
11.565°N
76.625°E
76.630°E
76.635°E
76.640°E
76.645°E
76.650°E
76.655°E
Masinagudi
800 m
6e+05
4e+05
2e+05
0e+00

Grey

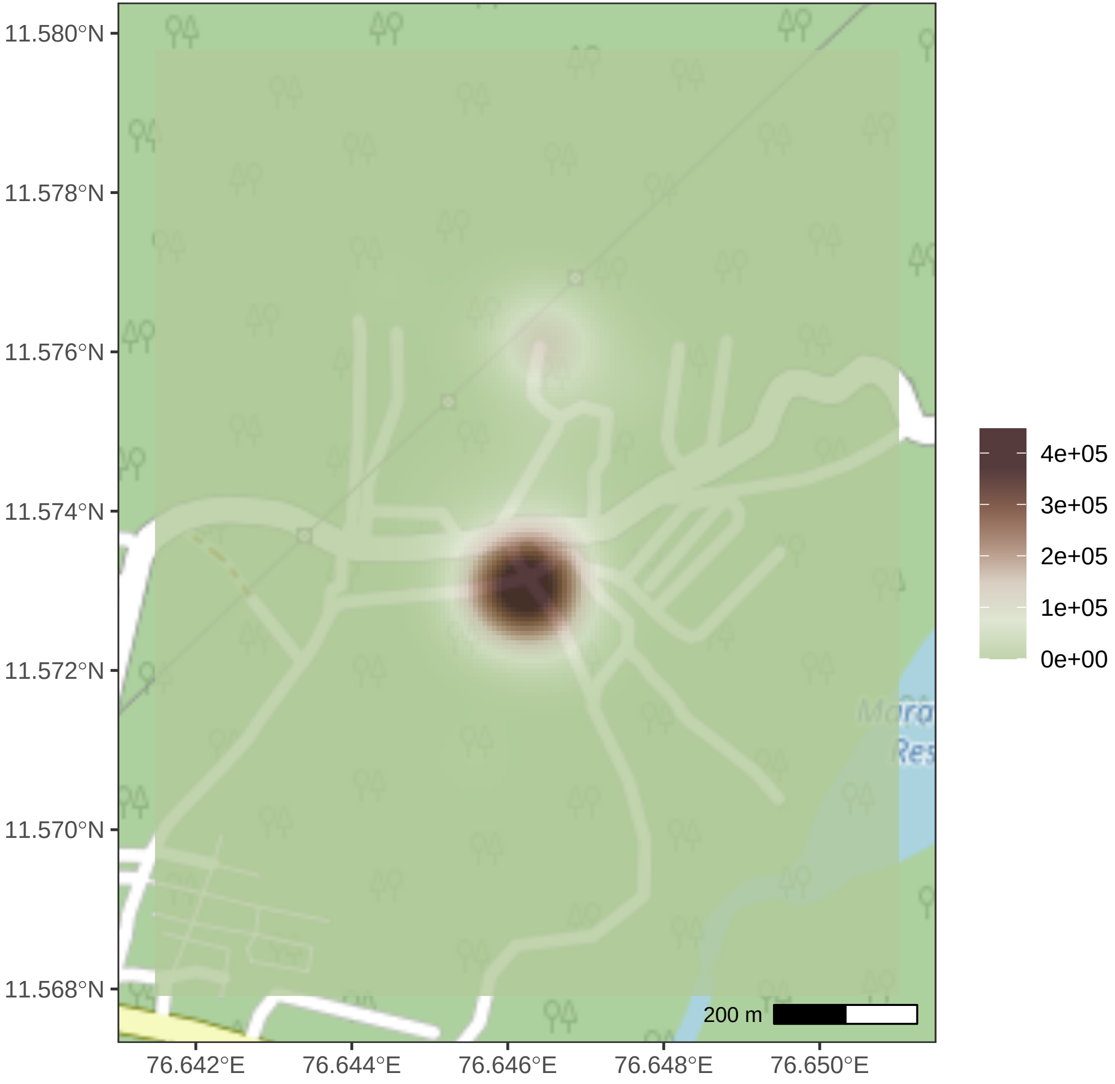
11.580°N
11.578°N
11.576°N
11.574°N
11.572°N
11.570°N
11.568°N
76.642°E
76.644°E
76.646°E
76.648°E
76.650°E
200 m
4e+05
3e+05
2e+05
1e+05
0e+00

Burger

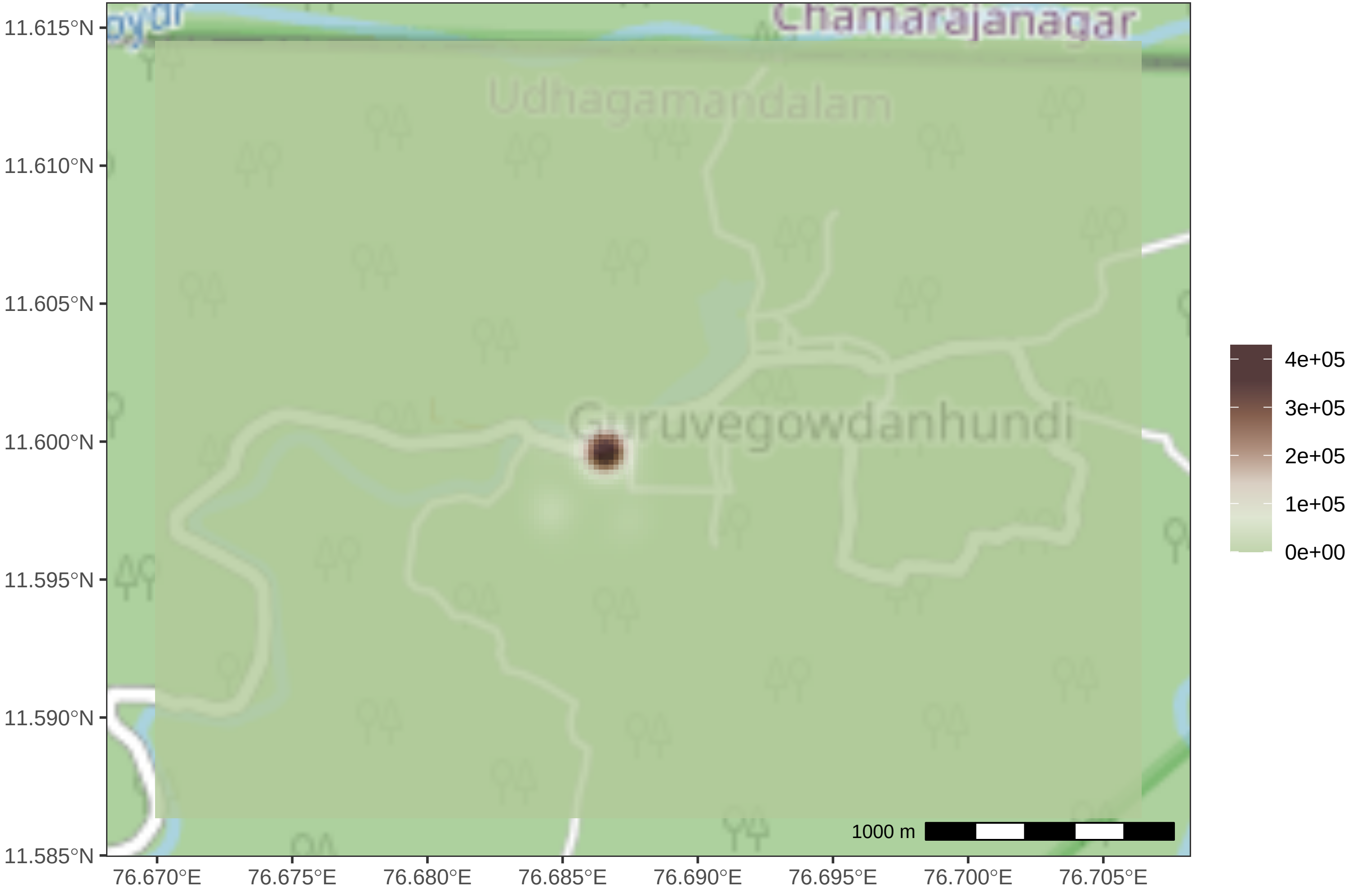

11.615°N
11.610°N
11.605°N
11.600°N
11.595°N
11.590°N
11.585°N
76.670°E
76.675°E
76.680°E
76.685°E
76.690°E
76.695°E
76.700°E
76.705°E
Chamarajanagar
Udhagamandalam
Guruvegowdanhundi
1000 m
4e+05
3e+05
2e+05
1e+05
0e+00

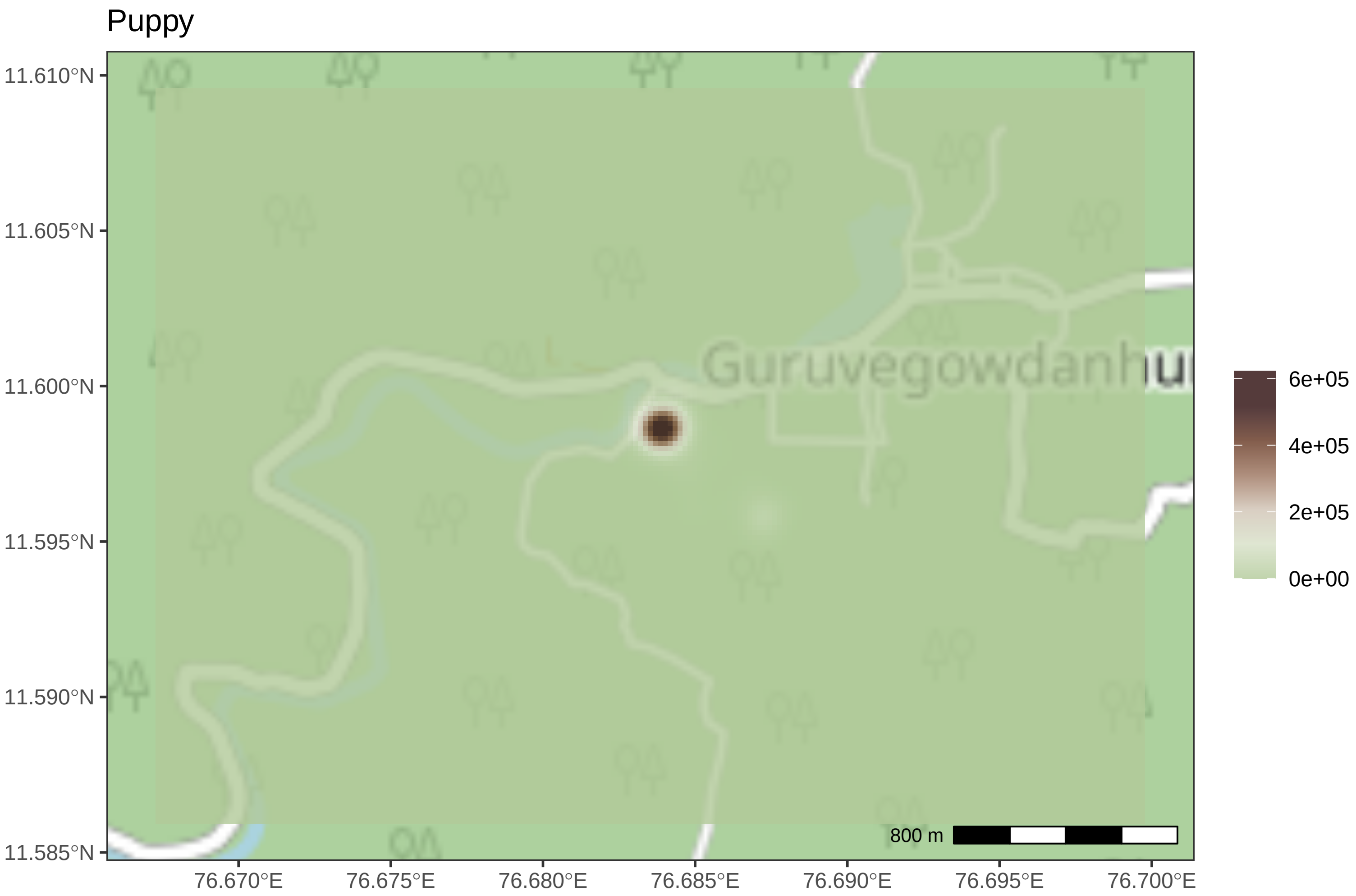

Puppy
11.610°N
11.605°N
11.600°N
11.595°N
11.590°N
11.585°N
76.670°E
76.675°E
76.680°E
76.685°E
76.690°E
76.695°E
76.700°E
Guruvegowdanhu
800 m
6e+05
4e+05
2e+05
0e+00

Snowy

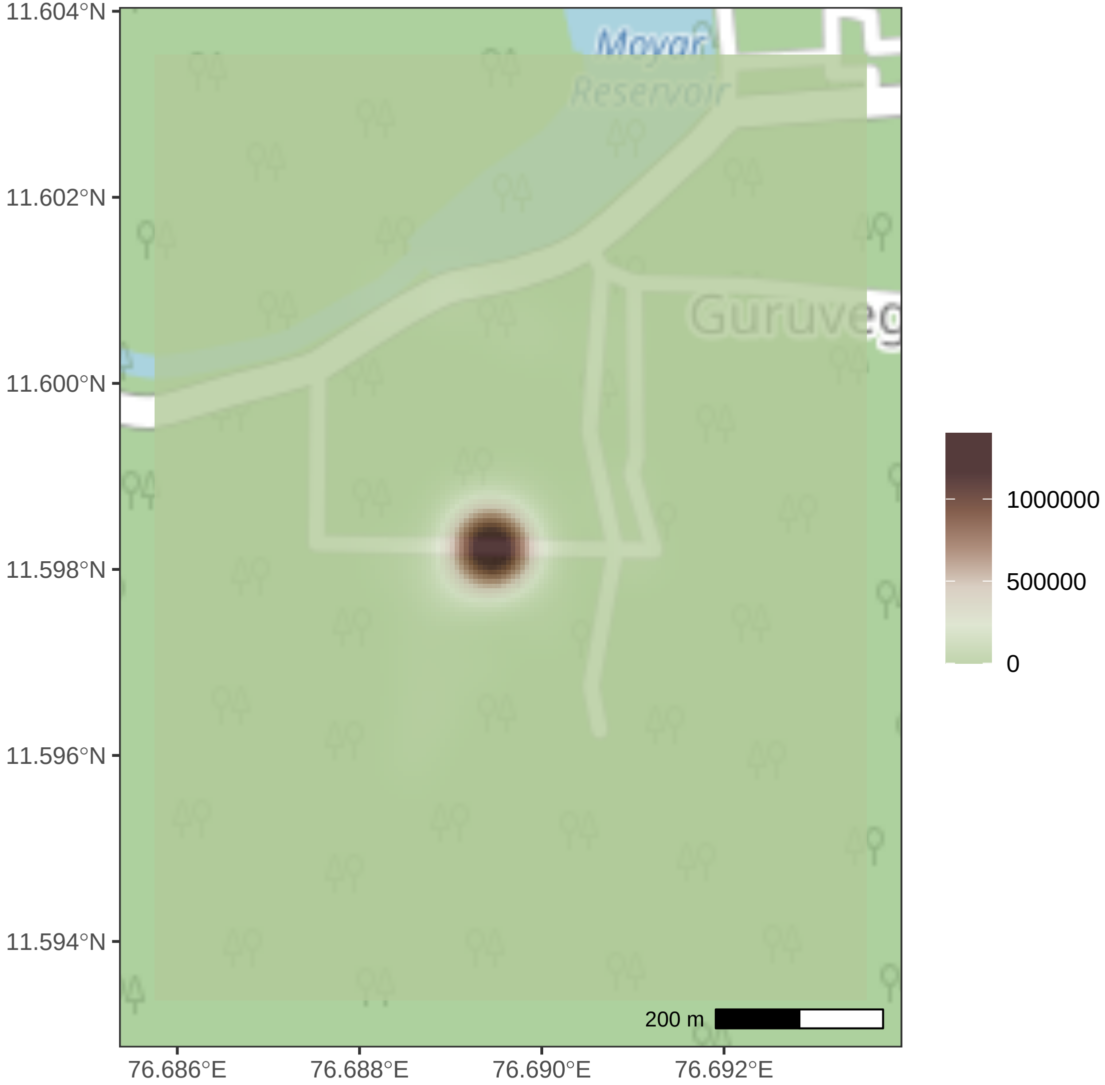
11.604°N
11.602°N
11.600°N
11.598°N
11.596°N
11.594°N
76.686°E
76.688°E
76.690°E
76.692°E
Moyar
Reservoir
200 m
1000000
500000
0

Chan

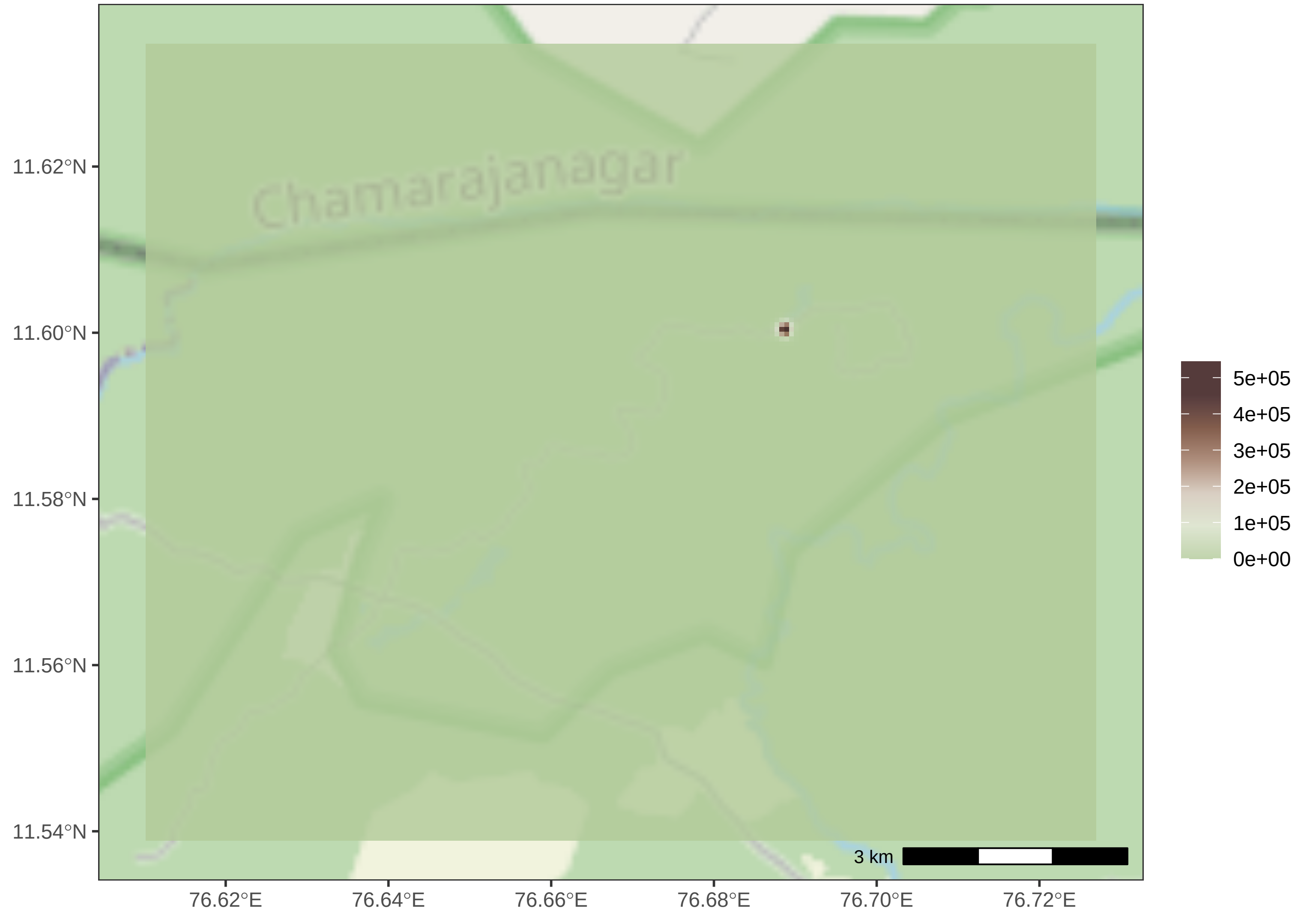
Chamarajanagar
11.62°N
11.60°N
11.58°N
11.56°N
11.54°N
76.62°E
76.64°E
76.66°E
76.68°E
76.70°E
76.72°E
3 km
5e+05
4e+05
3e+05
2e+05
1e+05
0e+00

Biscuit

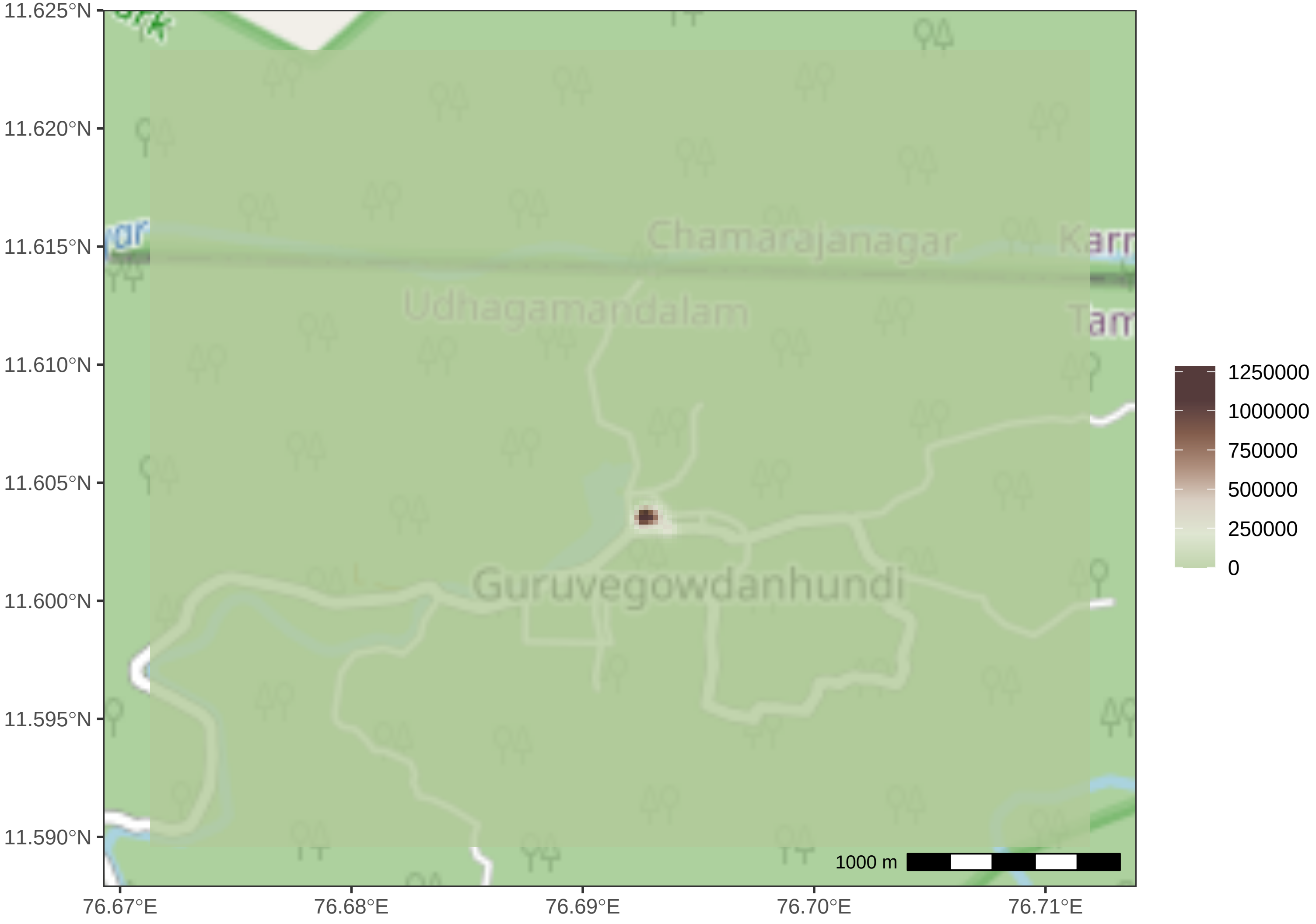

11.625°N
11.620°N
11.615°N
11.610°N
11.605°N
11.600°N
11.595°N
11.590°N
76.67°E
76.68°E
76.69°E
76.70°E
76.71°E
Chamarajanagar
Udhagamandalam
Guruvegowdanhundi
1000 m
1250000
1000000
750000
500000
250000
0

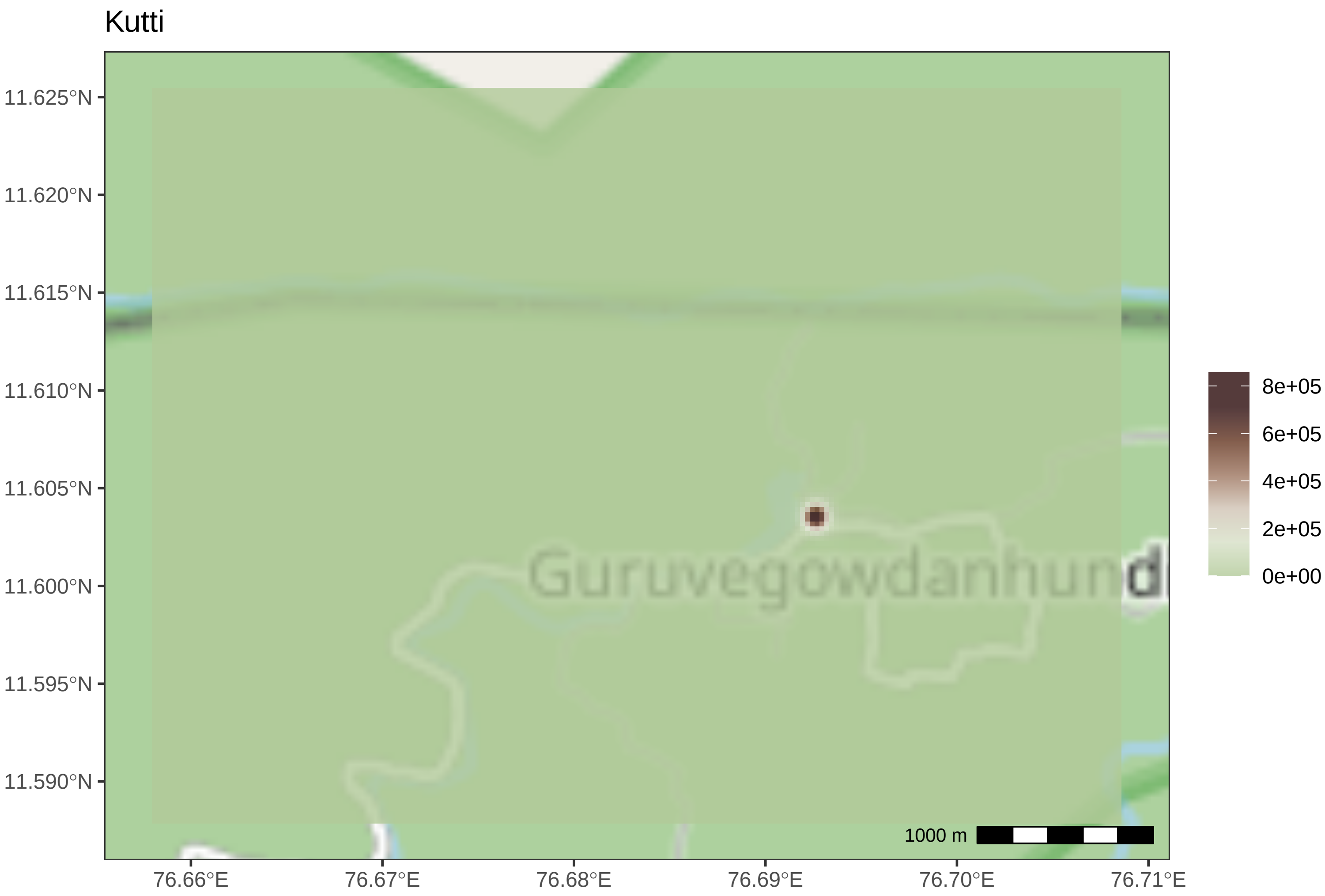
Kutti
11.625°N
11.620°N
11.615°N
11.610°N
11.605°N
11.600°N
11.595°N
11.590°N
76.66°E
76.67°E
76.68°E
76.69°E
76.70°E
76.71°E
Guruvegowdanhund
1000 m
8e+05
6e+05
4e+05
2e+05
0e+00

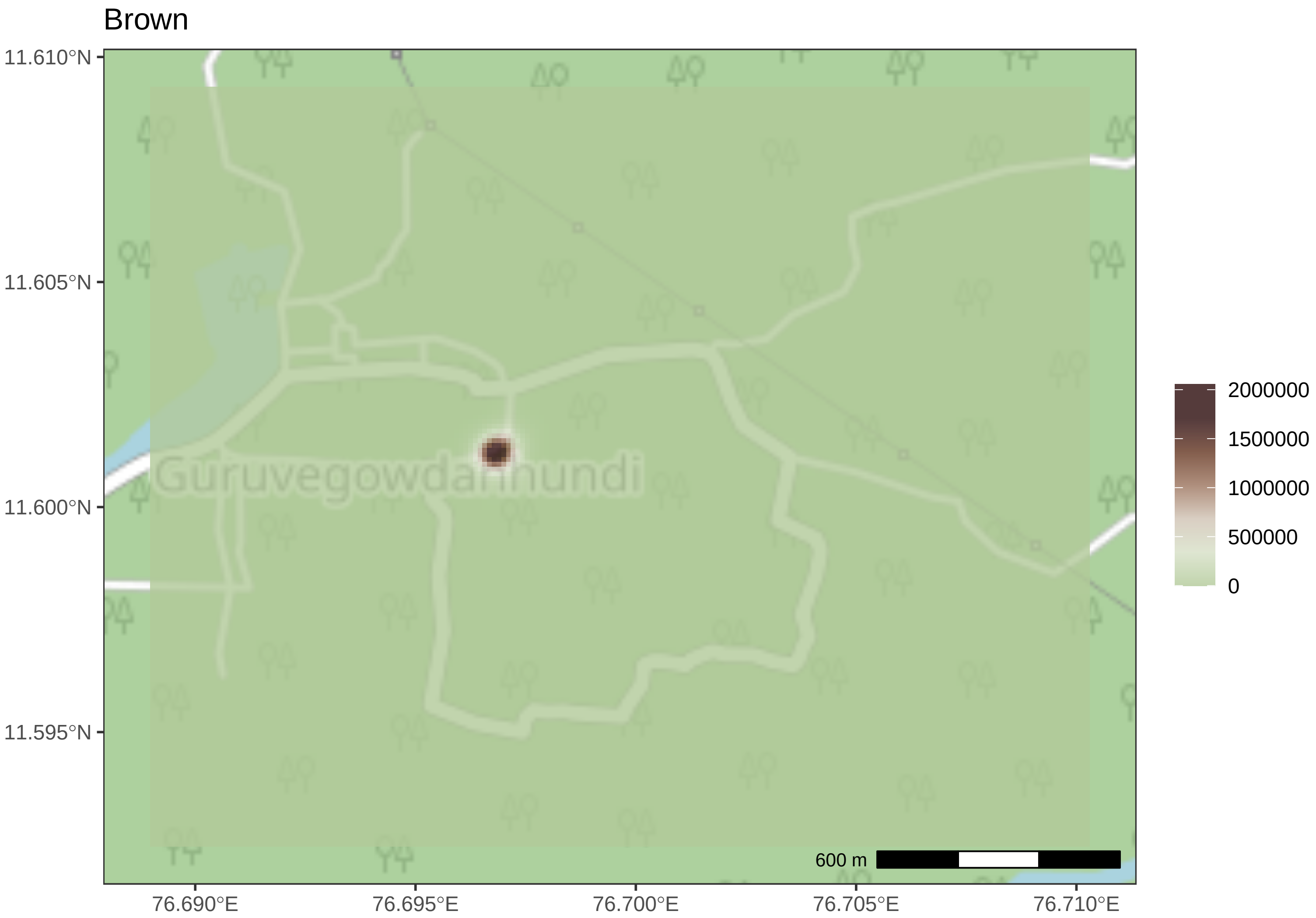
Brown
11.610°N
11.605°N
11.600°N
11.595°N
76.690°E
76.695°E
76.700°E
76.705°E
76.710°E
Guruvegowdanhundi
2000000
1500000
1000000
500000
0
600 m

Vellian

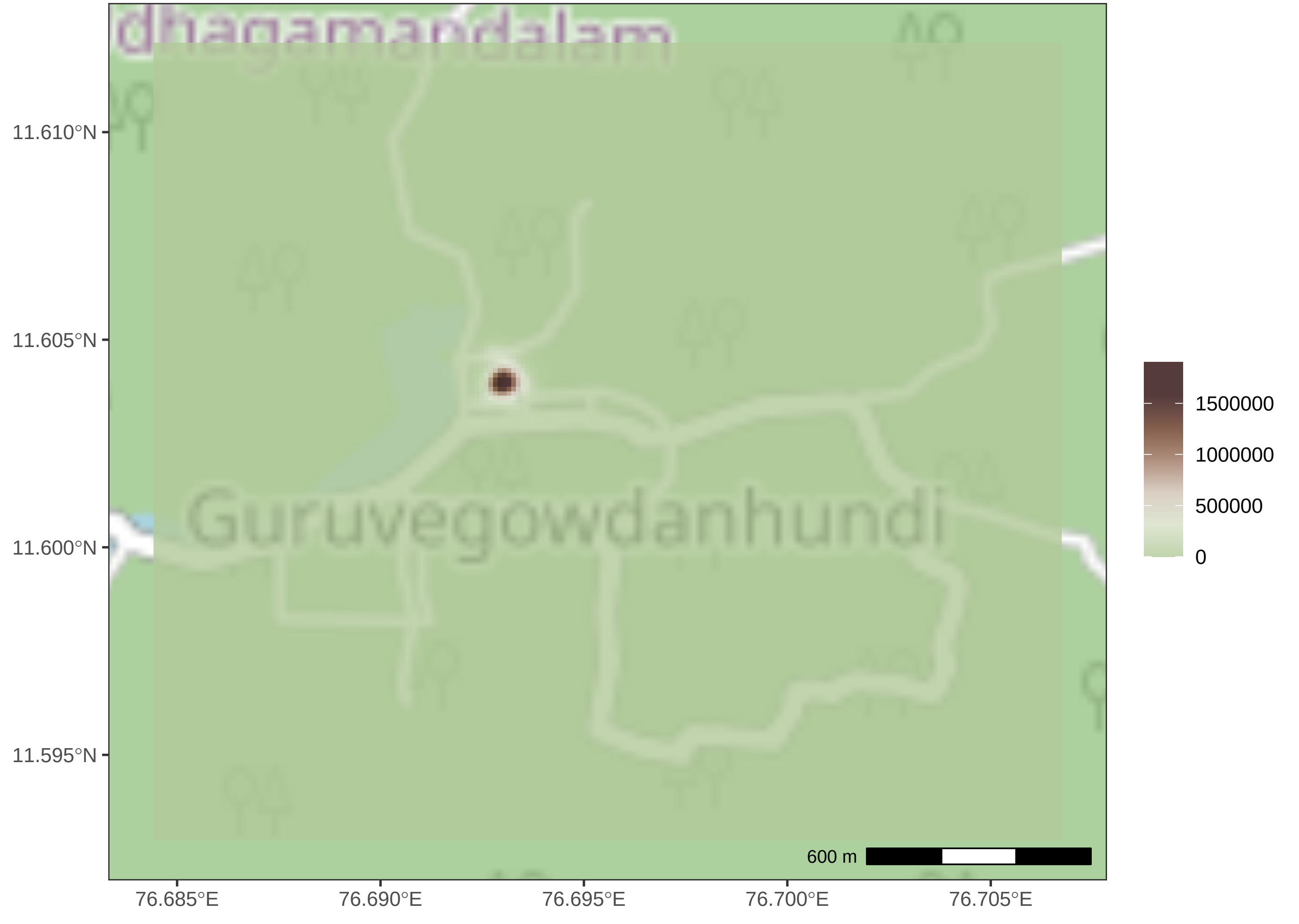

dhagamandalam
Guruvegowdanhundi
11.610°N
11.605°N
11.600°N
11.595°N
76.685°E
76.690°E
76.695°E
76.700°E
76.705°E
600 m
1500000
1000000
500000
0

PuppyMS

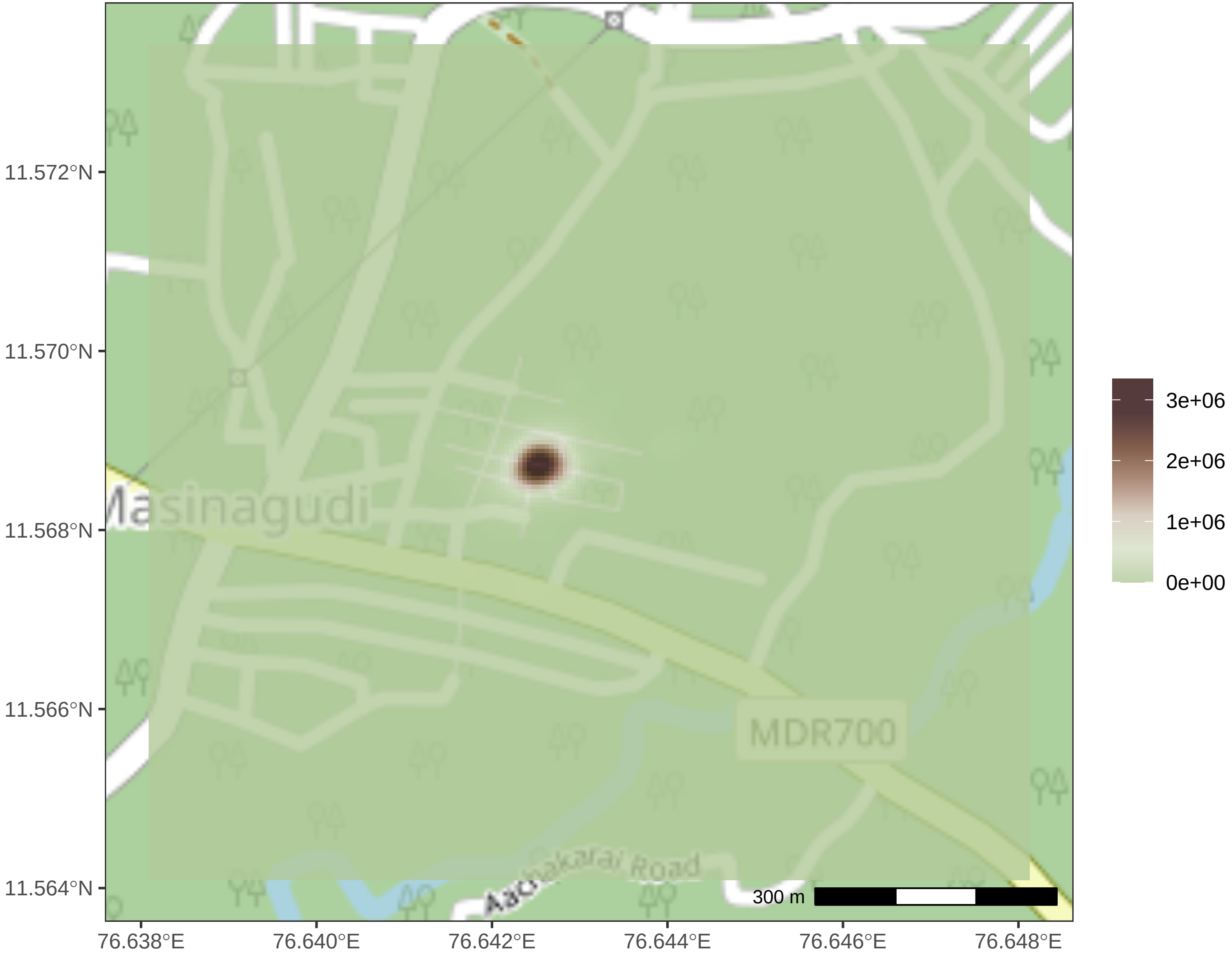
11.572°N
11.570°N
11.568°N
11.566°N
11.564°N
76.638°E
76.640°E
76.642°E
76.644°E
76.646°E
76.648°E
Masinagudi
MDR700
Road
300 m
3e+06
2e+06
1e+06
0e+00

Tony

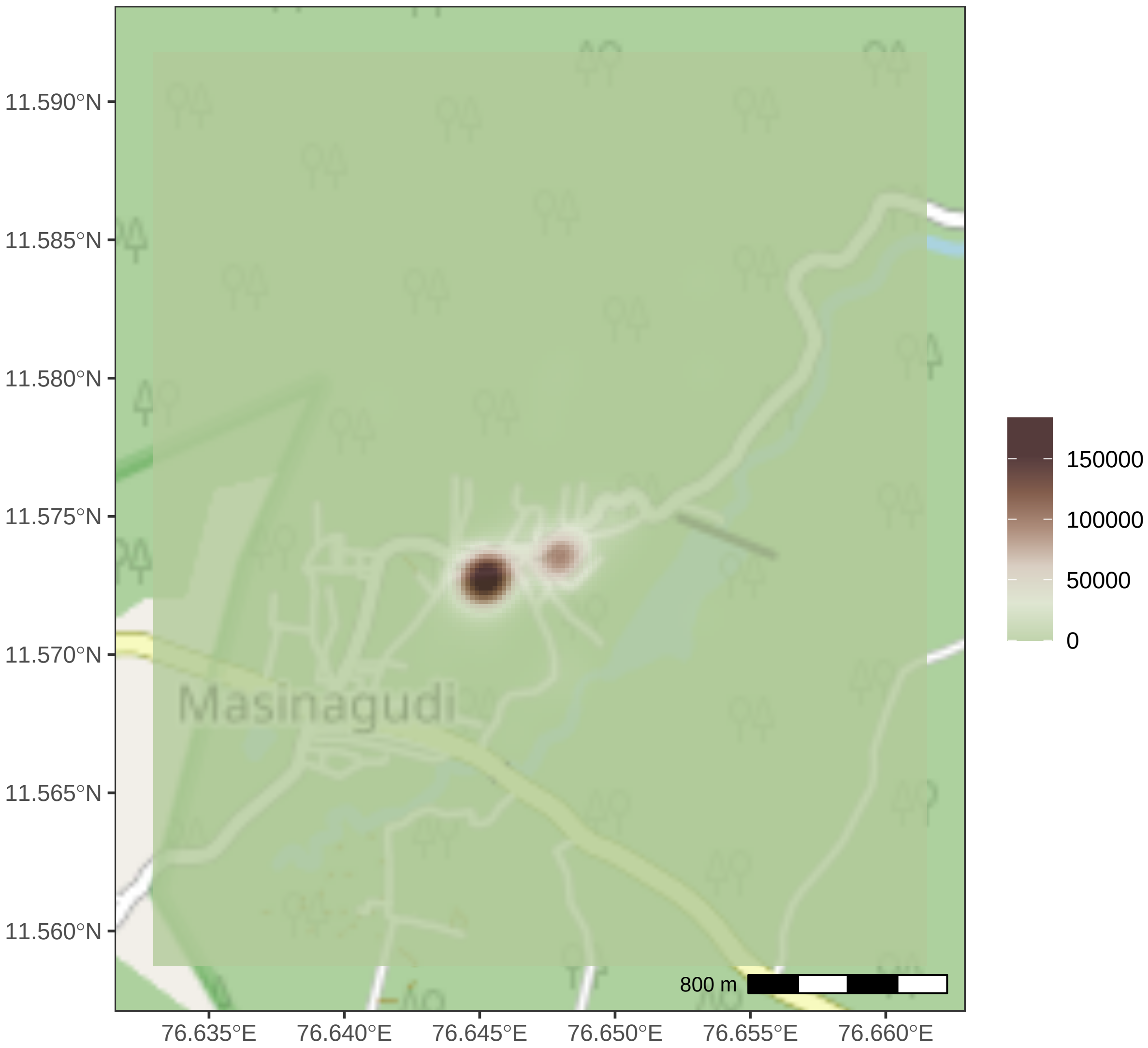

11.590°N
11.585°N
11.580°N
11.575°N
11.570°N
11.565°N
11.560°N
76.635°E
76.640°E
76.645°E
76.650°E
76.655°E
76.660°E
Masinagudi
800 m
150000
100000
50000
0

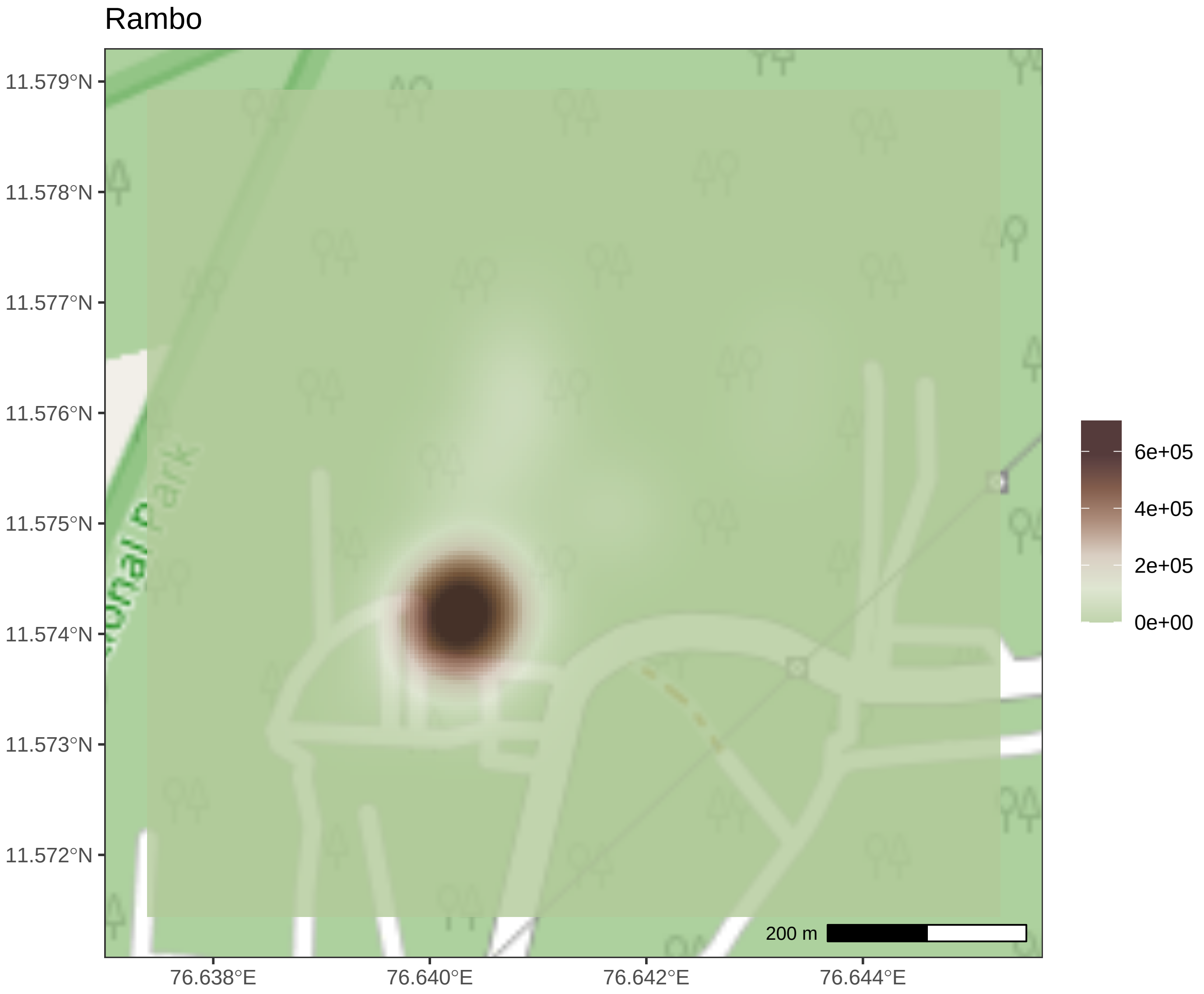
Rambo
11.579°N
11.578°N
11.577°N
11.576°N
11.575°N
11.574°N
11.573°N
11.572°N
76.638°E
76.640°E
76.642°E
76.644°E
6e+05
4e+05
2e+05
0e+00
200 m

Simba

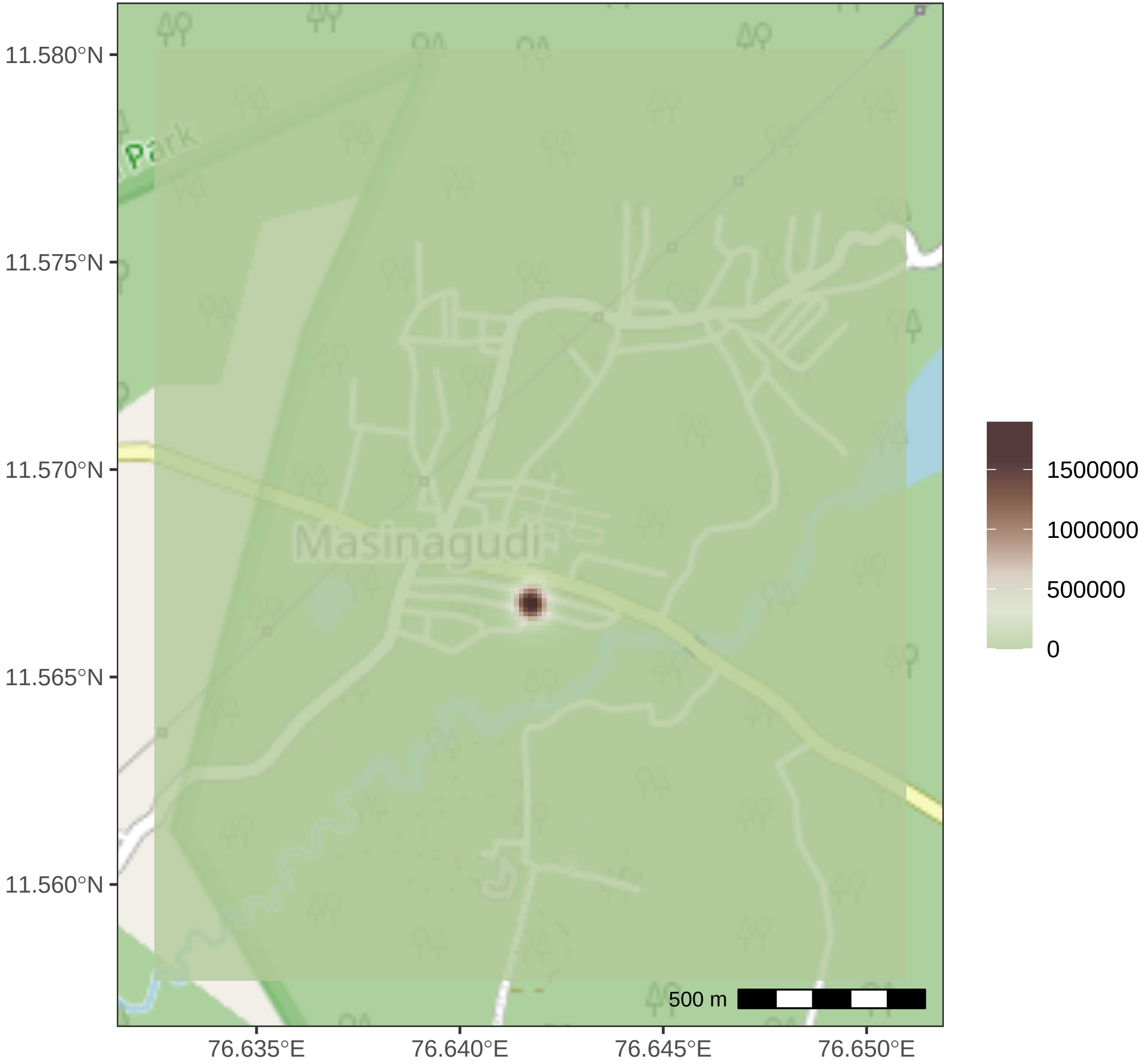

11.580°N
11.575°N
11.570°N
11.565°N
11.560°N
76.635°E
76.640°E
76.645°E
76.650°E
Masinagudi
Park
500 m
1500000
1000000
500000
0

### Meenu

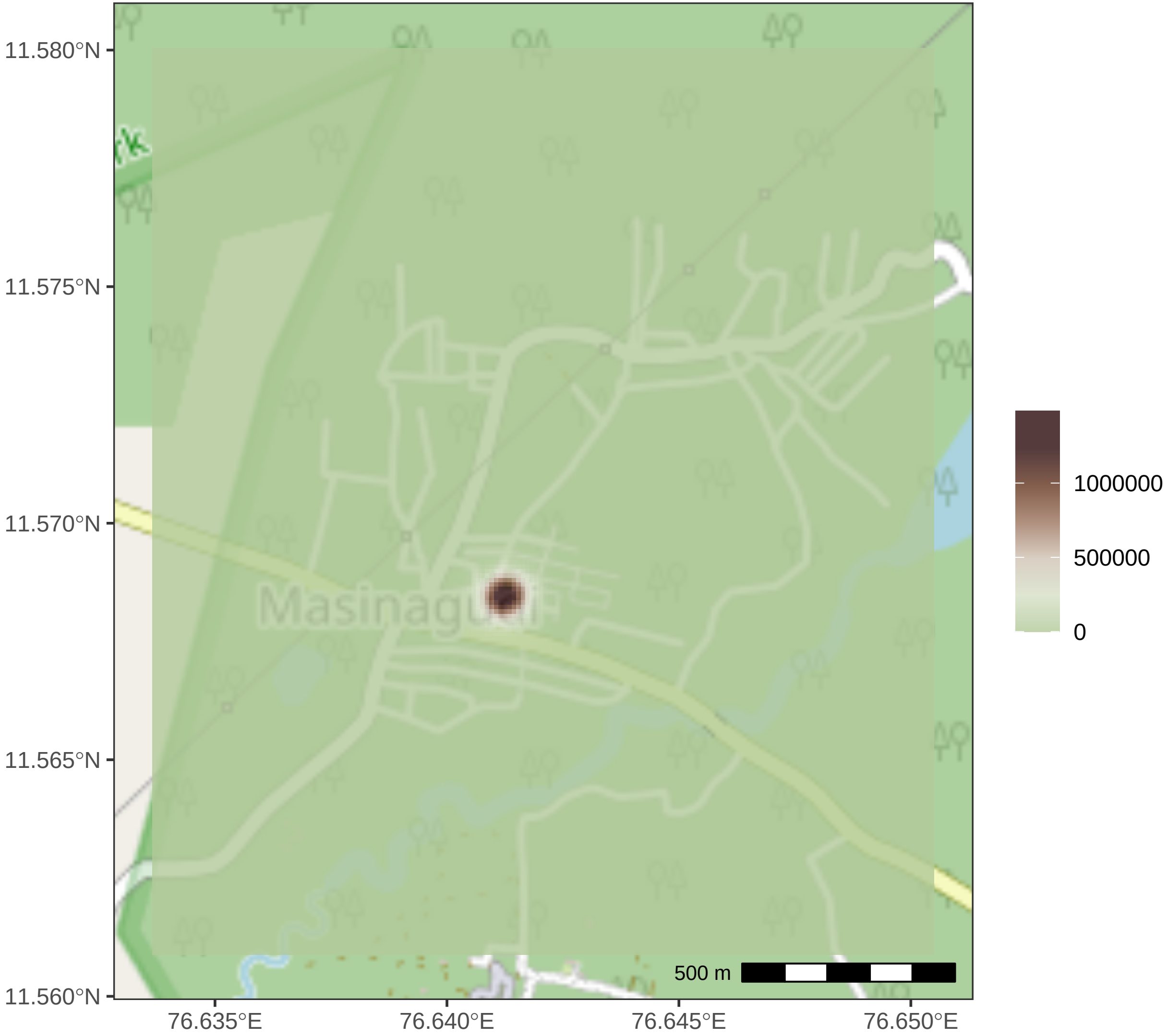
11.580°N
11.575°N
11.570°N
11.565°N
11.560°N
76.635°E
76.640°E
76.645°E
76.650°E
Masinagudi
500 m
1000000
500000
0

Blackie

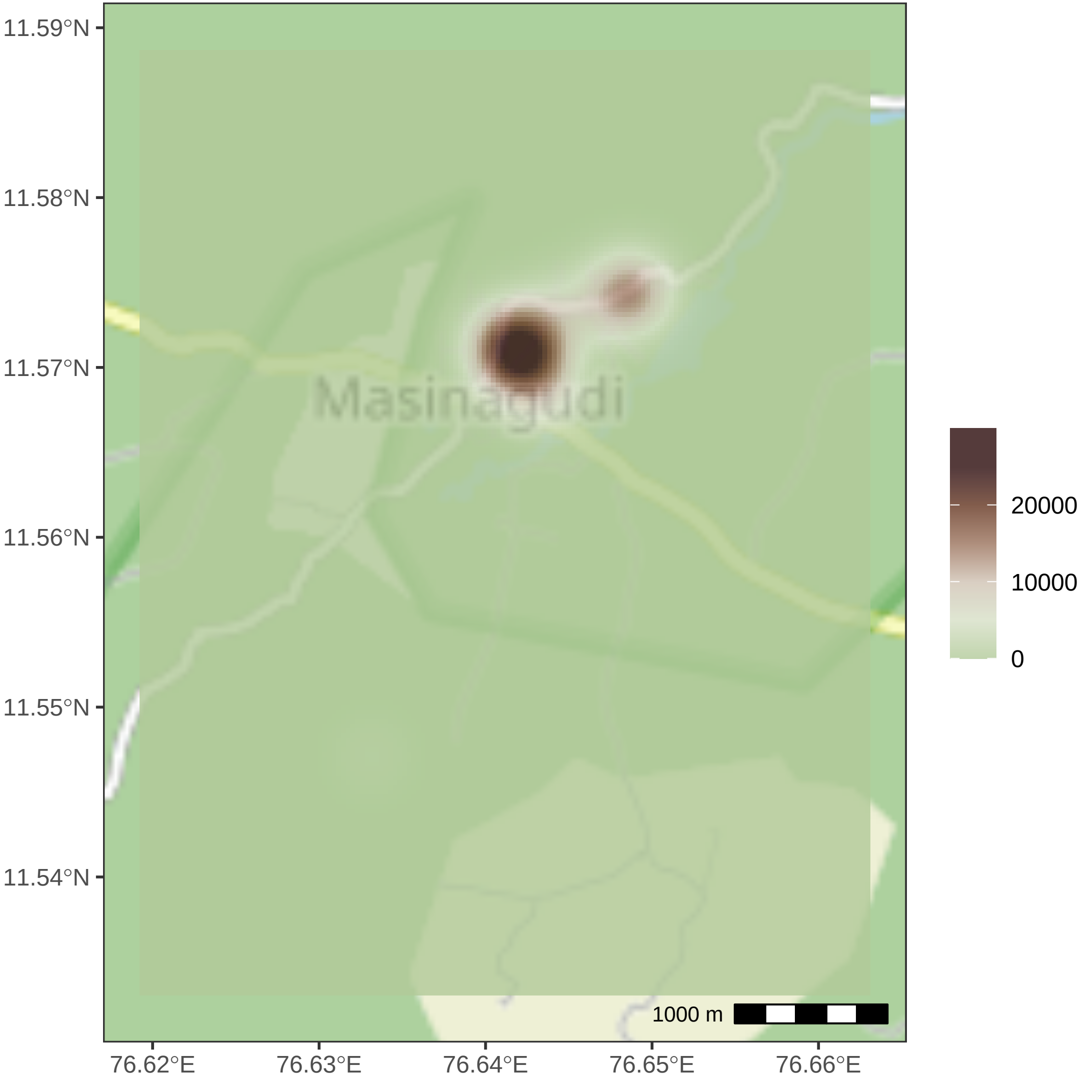

11.59°N
11.58°N
11.57°N
11.56°N
11.55°N
11.54°N
Masinagudi
1000 m
76.62°E
76.63°E
76.64°E
76.65°E
76.66°E
20000
10000
0

### Appu

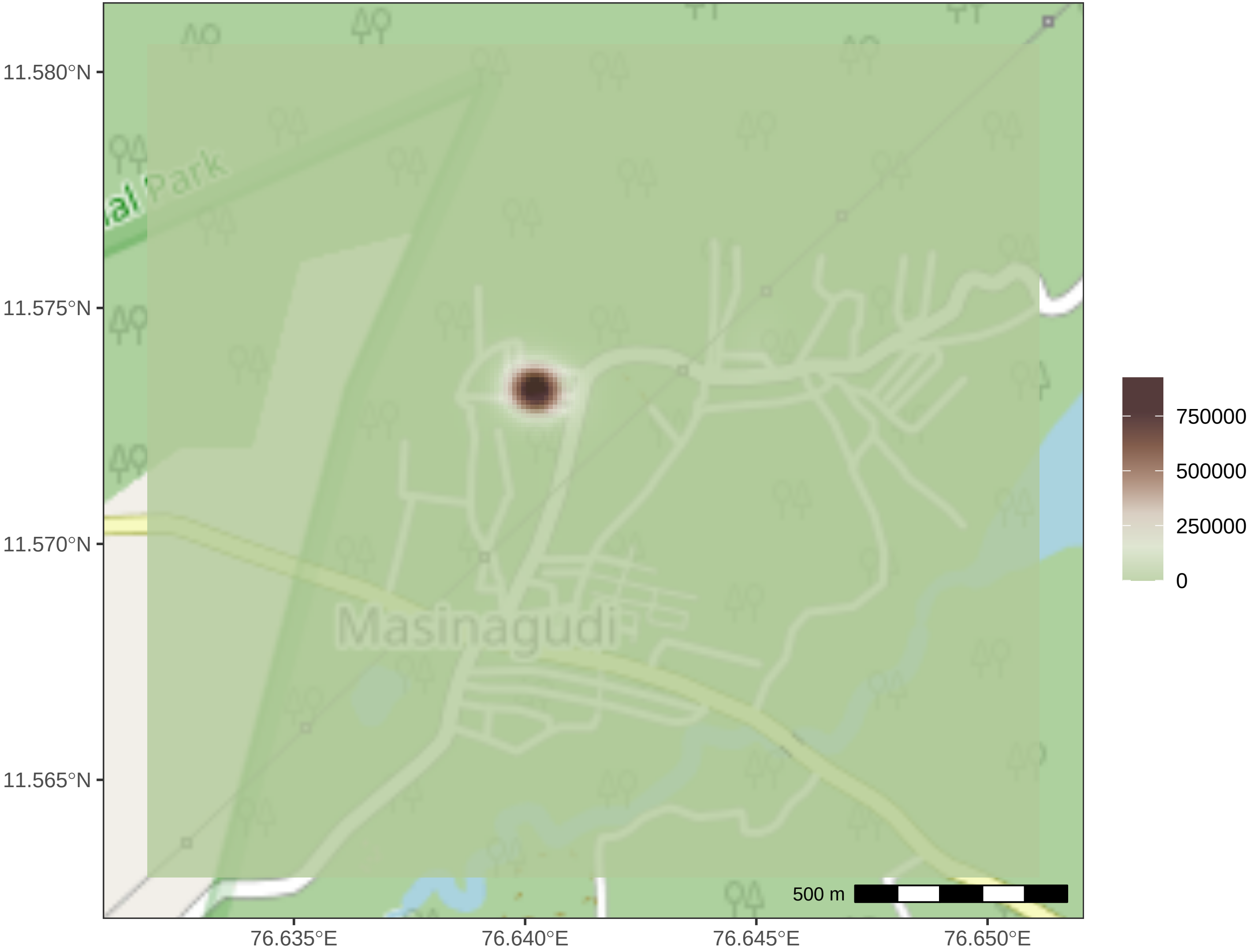

11.580°N
11.575°N
11.570°N
11.565°N
76.635°E
76.640°E
76.645°E
76.650°E
Park
Masinagudi
500 m
750000
500000
250000
0
